# Signals of polygenic adaptation on height have been overestimated due to uncorrected population structure in genome-wide association studies

**DOI:** 10.1101/355057

**Authors:** Mashaal Sohail, Robert M. Maier, Andrea Ganna, Alex Bloemendal, Alicia R. Martin, Michael C. Turchin, Charleston W. K. Chiang, Joel N. Hirschhorn, Mark J. Daly, Nick Patterson, Benjamin M. Neale, Iain Mathieson, David Reich, Shamil R. Sunyaev

## Abstract

Genetic predictions of height differ among human populations and these differences are too large to be explained by genetic drift. This observation has been interpreted as evidence of polygenic adaptation. Differences across populations were detected using SNPs genome-wide significantly associated with height, and many studies also found that the signals grew stronger when large numbers of subsignificant SNPs were analyzed. This has led to excitement about the prospect of analyzing large fractions of the genome to detect subtle signals of selection and claims of polygenic adaptation for multiple traits. Polygenic adaptation studies of height have been based on SNP effect size measurements in the GIANT Consortium meta-analysis. Here we repeat the height analyses in the UK Biobank, a much more homogeneously designed study. Our results show that polygenic adaptation signals based on large numbers of SNPs below genome-wide significance are extremely sensitive to biases due to uncorrected population structure.

## Introduction

Most human complex traits are highly polygenic.[1,2] For example, height has been estimated to be modulated by as much as 4% of human allelic variation.[2],[3] Polygenic traits are expected to evolve differently from monogenic ones, through slight but coordinated shifts in the frequencies of a large number of alleles, each with mostly small effect. In recent years, multiple methods have sought to detect selection on polygenic traits by evaluating whether shifts in the frequency of trait-associated alleles are correlated with the signed effects of the alleles estimated by genome-wide association studies (GWAS).[4–10]

Here we focus on a series of recent studies—some involving co-authors of the present manuscript—that have reported evidence of polygenic adaptation at alleles associated with height in Europeans. One set of studies observed that height-increasing alleles are systematically elevated in frequency in northern compared to southern European populations, a result that has subsequently been extended to ancient DNA.[4–11]Another study using a very different methodology (singleton density scores, SDS) found that height-increasing alleles have systematically more recent coalescent times in the United Kingdom (UK) consistent with selection for increased height over the last few thousand years.[12]

All of these studies have been based on SNP associations, in most cases with effect sizes discovered by the GIANT Consortium, which most recently combined 79 individual GWAS through meta-analysis, encompassing a total of 253,288 individuals.[13,14] Here, we show that the selection effects described in these studies are severely attenuated and in some cases no longer significant when using summary statistics derived from the UK Biobank, an independent and larger single study that includes 336,474 genetically unrelated individuals who derive their ancestry almost entirely from British Isles (identified as “white British ancestry” by the UK Biobank) (**Supplementary Table S1**). The UK Biobank analysis is based on a single cohort drawn from a relatively homogeneous population enabling excellent control of potential population stratification. Our analysis of the UK Biobank data confirms that almost all genome-wide significant loci discovered by the GIANT consortium are real associations, and that the two datasets have high concordance for low P value SNPs which do not reach genome-wide significance (**Supplementary Figure S1**; genetic correlation between the two height studies is 0.94 [se=0.0078]). However, our analysis yields qualitatively different conclusions with respect to signals of polygenic adaptation.

## Results

We began by estimating “polygenic height scores”—sums of allele frequencies at independent SNPs from GIANT weighted by their effect sizes—to study population level differences among ancient and present-day European samples. We used a set of different significance thresholds and strategies to correct for linkage disequilibrium as employed by previous studies, and replicated their signals for significant differences in genetic height across populations.[4–11] (**Figure 1a, Supplementary Figure S2**). We then repeated the analysis using summary statistics from a GWAS for height in the UK Biobank restricting to individuals of British Isles ancestry and correcting for population stratification based on the first ten principal components (UKB).[15] This analysis resulted in a dramatic attenuation of differences in polygenic height scores (**Figure 1a, Supplementary Figures S2–S4**). The differences between ancient European populations also greatly attenuated (**Figure 1a, Supplementary Figure S5**). Strikingly, the ordering of the scores for populations also changed depending on which GWAS was used to estimate genetic height both within Europe (**Figure 1a, Supplementary Figures S2–S5**) and globally (**Supplementary Figure S6**), consistent with reports from a recent simulation study.[16] The height scores were qualitatively similar only when we restricted to independent genome-wide significant SNPs in GIANT and the UK Biobank (P < 5×10^−8^) (**Supplementary Figure S2b**). This replicates the originally reported significant north-south difference in the allele frequency of the height-increasing allele[4] or in genetic height[5] across Europe, as well as the finding of greater genetic height in ancient European steppe pastoralists than in ancient European farmers,[6] although the signals are attenuated even here. This suggests that tests of polygenic adaptation based on genome-wide significant SNPs may be relatively insensitive to confounding (**Supplementary Figure S2b**), and that confounding due to stratification is a particular danger for sub-significant SNPs (**Figure 1a, Supplementary Figure S2a**).

**Figure 1.**
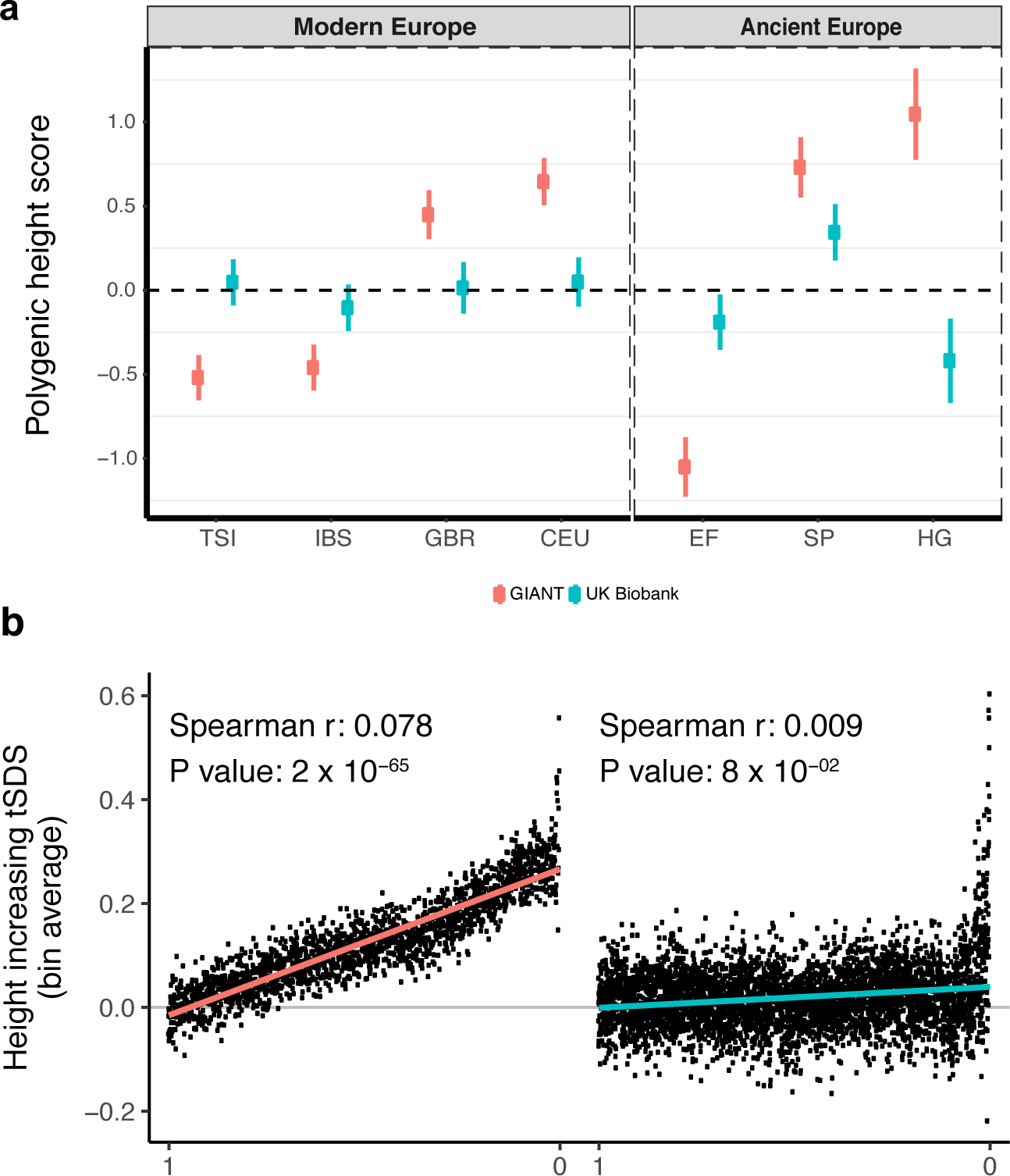
Polygenic height scores and tSDS scores based on GIANT and UK Biobank GWAS. Polygenic scores in present-day and ancient European populations are shown, centered by the average score across populations and standardized by the square root of the additive variance. Independent SNPs for the polygenic score from both GIANT (*red*) and the UK Biobank ***(blue)*** were selected by picking the SNP with the lowest P value in each of 1700 independent LD blocks similarly to refs [8,9] (see methods). Present-day populations are shown from Northern Europe (CEU, GBR) and Southern Europe (IBS, TSI) from the 1000 genomes project; Ancient populations are shown in three meta-populations (HG = Hunter-Gatherer (n=162 individuals), EF = Early Farmer (n=485 individuals), and SP = Steppe Ancestry (n=465 individuals)) (see **Supplementary Table S2**). Error bars are drawn at 95% credible intervals. See **Supplementary figures S2–S6** for polygenic height scores computed using other linkage disequilibrium pruning procedures, significance thresholds, summary statistics and populations. (b) tSDS for height-increasing allele in GIANT (left) and UK Biobank (right). The tSDS method was applied using pre-computed Singleton Density Scores for 4,451,435 autosomal SNPs obtained from 3,195 individuals from the UK10K project[12]’[17] for SNPs associated with height in GIANT and the UK biobank. SNPs were ordered by GWAS P value and grouped into bins of 1000 SNPs each. The mean tSDS score within each P value bin is shown on the y-axis. The Spearman correlation coefficient between the tSDS scores and GWAS P values, as well as the correlation standard errors and P values, were computed on the un-binned data. The gray line indicates the null-expectation, and the colored lines are the linear regression fit. The correlation is significant for GIANT (Spearman r = 0.078, P = 1.55 x 10^−65^) but not for UK Biobank (Spearman r = −0.009, P = 0.077).

Next, we looked at polygenic adaptation within the UK using the “singleton density score” (SDS)—an independent measure that uses the local density of alleles that occur only once in the sample as a proxy for coalescent branch lengths.[12,17] SDS can be combined with GWAS effect sizes estimates by aligning the SDS sign to the trait-increasing allele, after which the score is referred to as tSDS. A tSDS score larger than zero implies that height-increasing alleles have been increasing in frequency over time due to natural selection. We replicate the finding that tSDS computed in the UK10K is positively rank-correlated with GIANT[12] height P values (Spearman’s ρ = 0.078, P = 1.55 x 10^−65^, **Figure 1b**). However, this signal of polygenic adaptation in the UK attenuated when we used UK Biobank height effect size estimates and P values and became formally non-significant (ρ = 0.009, P = 0.077, **Figure 1b**).

We propose that the qualitative difference between the polygenic adaptation signals in GIANT and the UK Biobank is the cumulative effect of subtle biases in each of the contributing SNPs in GIANT. This bias can arise due to incomplete control of the population structure in GWAS.[18] For example, if height were differentiated along a north-south axis because of differences in environment, any variant that is differentiated in frequency along the same axis would have an artifactually large effect size estimated in the GWAS. Population structure is substantially less well controlled for in the GIANT study than in the UK Biobank study, both because the GIANT study population is more heterogeneous than that in the UK Biobank, and because the population structure in GIANT may not have been well controlled in some component cohorts of GIANT due to the relatively small size of individual studies (i.e., the ability to detect and correct population structure is dependent on sample size[19,20]). The GIANT meta-analysis also found that such stratification effects worsen as SNPs below genome-wide significance are used to estimate height scores,[14] consistent with our finding that the differences in genetic height increase when including these SNPs.

To obtain further insight into our observed discrepancy between polygenic adaptation signals in GIANT vs the UK Biobank, we repeated our analyses using estimates of height effect sizes computed using different methods, and then interrogated each of these for signs of population structure. Repeating our analysis with family-based effect size estimates from an independent study (NG2015 sibs),[7] we found evidence for significant differences in polygenic scores between northern and southern Europeans that were qualitatively similar to those obtained using GIANT effect size estimates (**Supplementary Figure S4–S5**). Inclusion of individuals from the UK Biobank who were not of British Isles ancestry without controlling for population structure (UKB all no PCs) in the measurements of effect sizes also produced this pattern (**Supplementary Figure S3–S5**). Thus, UK Biobank estimates that retain population structure show similar patterns to GIANT and previously published family-based estimates (NG 2015 sibs). In contrast, no significant signals of genetic stratification of height or a strong tSDS signal are present across populations from: 1) a genetically homogeneous sample of UK Biobank with entirely British Isles ancestry without controlling for population structure (UKB WB no PCs), or 2) effect size estimates based on UK Biobank families (UKB sibs, UKB sibs WB) (**Supplementary Figures S3–S5, S7–S8**). These analyses provide further evidence that the lack of signal in the UK Biobank analysis is unlikely to be simply due to over-correction for structure in the original UKB estimates.

We obtained direct confirmation that population structure is more correlated with effect size estimates in GIANT than to those in the UK Biobank. **Figure 2a** shows that the effect sizes estimated in GIANT are highly correlated with the SNP loadings of several principal components of population structure (PC loadings). Previously published family-based effect size estimates[7] (NG 2015 sibs) are similarly correlated with the PC loadings showing that they are also affected by population structure despite being computed within families; in other words, these empirical analyses show that even previously published family-based effect size estimates are not free from concerns about population structure. The within-family strategy for eliminating concerns about population stratification is not problematic on its own as our UK Biobank family estimates (UKB sibs, UKB sibs WB) computed using the same method do not show any stratification effects (**Supplementary Figures S10–S12**). We also do not see a strong correlation with PC loadings in our UK Biobank estimates computed using unrelated individuals (UKB)(**Figure 2a**). However, the UK Biobank estimates including individuals not of British Isles ancestry and not correcting for population structure (UKB all no PCs) show the same stratification effects as GIANT and NG2015 sibs (**Supplementary Figure S10–S12**). Similarly, we find that alleles that are more common in the Great Britain population (GBR) than in the Tuscan population from Italy (TSI) tend to be preferentially height-increasing according to the GIANT and NG2015 sibs estimates but not according to the UKB estimates (**Figure 2c, Supplementary Figures S11, S12**).

**Figure 2.**
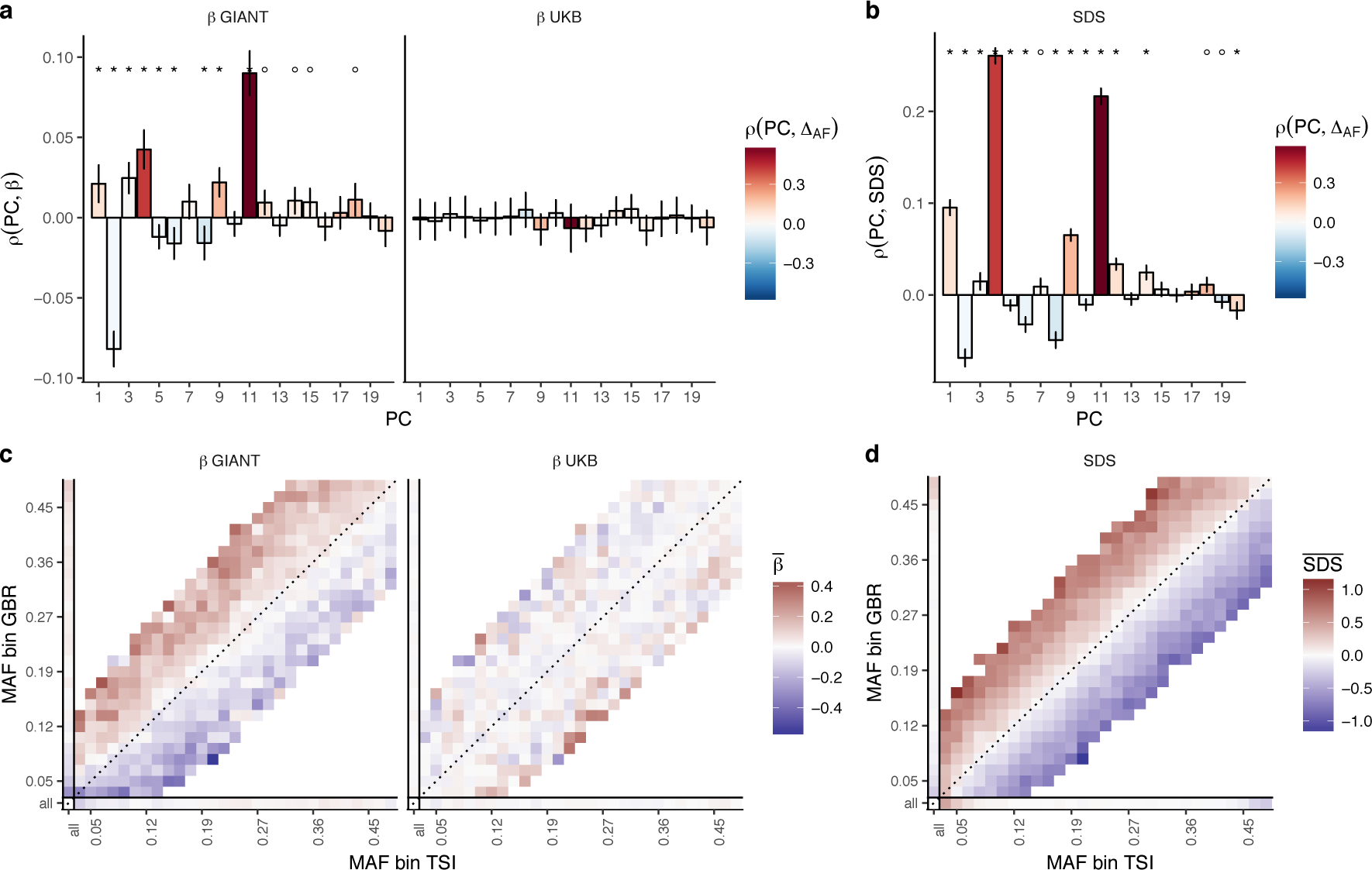
Evidence of stratification in height summary statistics. Top row: Pearson Correlation coefficients of (a) PC loadings and height beta coefficients from GIANT and UKB, and (b) PC loadings and SDS (pre-computed in the UK10K) across all SNPs. PCs were computed in all 1000 genomes phase 1 samples. Colors indicate the correlation of each PC loading with the allele frequency difference between GBR and TSI, a proxy for the European North-South genetic differentiation. PC 4 and 11 are most highly correlated with the GBR - TSI allele frequency difference. Confidence intervals and P values are based on Jackknife standard errors (1000 blocks). Open circles indicate correlations significant at alpha = 0.05, stars indicate correlations significant after Bonferroni correction in 20 PCs (P < 0.0025). Bottom row: Heat map after binning all SNPs by GBR and TSI minor allele frequency of (c) mean beta coefficients from GIANT and UKB, and (d) SDS scores for all SNPs. Only bins with at least 300 SNPs are shown. While the stratification effect in SDS is not unexpected, it can lead to false conclusions when applied to summary statistics that exhibit similar stratification effects. UKB height betas exhibit stratification effects that are weaker, and in the opposite direction of the stratification effects in GIANT (see **Supplementary Figure S9** for a possible explanation).

The tSDS analysis should be robust to the type of population structure discussed above.[12] However, there is a north-south cline in singleton density in Europe due to the lower genetic diversity in northern than in southern Europeans, with singleton density being lower in northern than in southern regions.[21] As a consequence, SDS tends to be higher in alleles more common in GBR than in TSI (**Figure 2d**). This cline in singleton density coincidentally parallels the phenotypic cline in height and the major axis of genome-wide genetic variation. Therefore, when we perform the tSDS test using GIANT-estimated effect sizes and P values, we find fewer singletons (corresponding to higher SDS) around the inferred height-increasing alleles which tend, due to the uncontrolled population stratification in GIANT, to be at high frequency in northern Europe (**Figure. 2c**). This effect does not appear when we use UK Biobank summary statistics because of the much lower level of population stratification and more modest variation in height. We find that SDS is not only correlated with GBR-TSI allele frequency differences, but with several principal component loadings across all SNPs (**Figure 2b**), and that these SDS-PC correlations often coincide with correlations between GIANT-estimated effect sizes and PC loadings (**Figure 2a**).

We further find that the tSDS signal which is observed across the whole range of P values in some summary statistics can be mimicked by replacing SDS with GBR-TSI allele frequency differences (**Figure. 3a, 3c, Supplementary Figures S7–S8, S13–S14**), suggesting that the tSDS signal at non-significant SNPs may be driven in part by residual population stratification. As with the polygenic score analysis, a small but significant effect is observed when we restrict to genome-wide significant SNPs (P < 5 x 10^−8^). This effect persists when using UK Biobank family-based estimates for genome-wide significant SNPs (**Figure 3b**), and is not driven by allele frequency differences between GBR and TSI (**Figure 3d**), suggesting a true but attenuated signal of polygenic adaptation in the UK that is driven by a much smaller number of SNPs than previously thought. Indeed, a tSDS signal which is driven by natural selection is not expected to lead to an almost linear increase over the whole P value range in a well-powered GWAS. Instead, we would expect to see a greater difference between highly significant SNPs and non-significant SNPs, similar to the pattern observed in the UK Biobank (**Figure 3a**).

**Figure 3.**
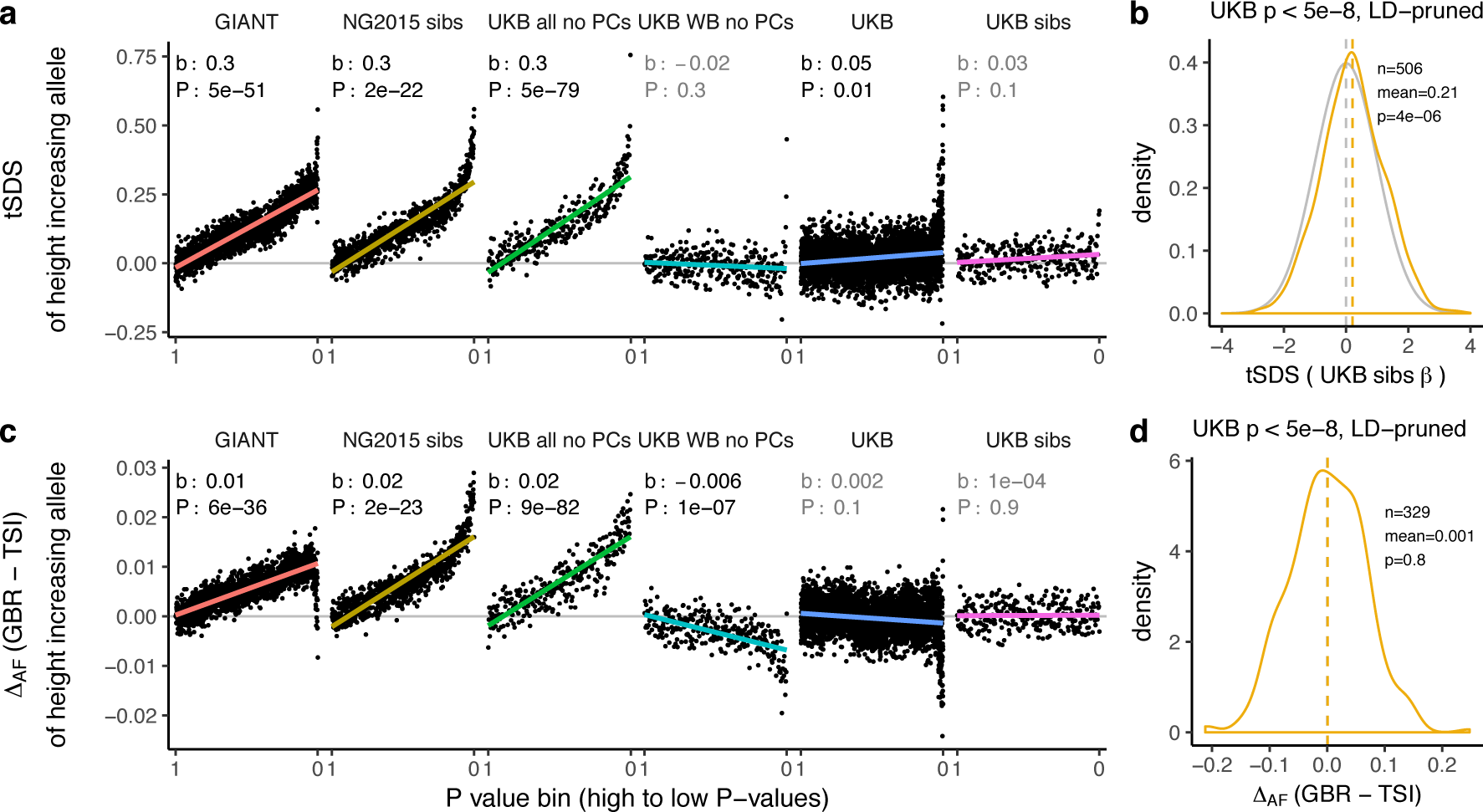
Height tSDS results for different summary statistics. Mean tSDS of the height increasing allele in each P value bin for six different summary statistics. The first two panels are computed analogously to Figure 4A and Figure S22 in Field et al. In contrast to those Figures and to **Figure 1b**, the displayed betas and P values correspond to the slope and P value of the linear regression across all un-binned SNPs (rather than the Spearman correlation coefficient and Jackknife P values). The y-axis has been truncated at 0.75, and does not show the top bin for UKB all no PCs, which has a mean tSDS of 1.5. See **Supplementary Figure S7** for other GWAS summary statistics (b) tSDS distribution of the height increasing allele in 506 LD-independent SNPs which are genome-wide significant in a UKB height GWAS, where the beta coefficient is taken from a within sibling analysis in the UKB. The gray curve represents the standard normal null distribution, and we observe a significant shift providing confirmation of a real SDS signal of polygenic adaptation for height. (c) Allele frequency difference between GBR and TSI of the height increasing allele in each P value bin for six different summary statistics. Betas and P values correspond to the slope and P value of the linear regression across all un-binned SNPs. The lowest P value bin in UKB all no PCs with a y-axis value of 0.06 has been omitted. See **Supplementary Figure S13** for other GWAS summary statistics. (d) Allele frequency difference between GBR and TSI of the height increasing allele in 329 LD-independent SNPs which are genome-wide significant in a UKB height GWAS and were intersecting with our set of 1000 genomes SNPs. There is no significant difference in frequency in these two populations, suggesting that tSDS shift at the gw-significant SNPs is not driven by population stratification. The patterns shown here suggest that the positive tSDS values across the whole range of P values is a consequence of residual stratification. At the same time, the increase in tSDS at genome-wide significant, LD-independent SNPs in (b) cannot be explained by GBR - TSI allele frequency differences as shown in (d). Binning SNPs by P value without LD-pruning can lead to unpredictable patterns at the low P value end, as the SNPs at the low P value end are less independent of each other than higher P value SNPs (**Supplementary Figure S15**). **Supplementary Figures S8 and S14** therefore show the same data for a set of LD-pruned SNPs.

Lastly, we asked whether any remaining differences in polygenic height scores among populations are driven by polygenic selection by using the Q_x_ framework to test against a null model of genetic drift.[5] We re-computed polygenic height scores in the POPRES dataset for this analysis as it has larger sample sizes of northern and southern Europeans than the 1000 Genomes project.[22] We computed height scores using independent SNPs that are 1) genome-wide significant in the UK Biobank (“gw-sig”, P < 5 x 10^−8^) and 2) sub-significantly associated with height (“sub-sig”, P < 0.01) in different GWAS datasets. For each of these, we tested if population differences were significant due to an overall overdispersion (Pqx), and if they were significant along a north-south cline (Pl at) (**Figure 4, Supplementary Figure S16–S17**). Both gw-sig and sub-sig SNP-based scores computed using GIANT effect sizes showed significant overdispersion of height scores overall and along a latitude cline, consistent with previous results (**Figure 4, Supplementary Figure S16–S17**). However, the signal attenuated dramatically between sub-sig (Q_x_ = 1100, P_Qx_ = 1 x 10^−220^) and gw-sig (Q_x_ = 48, P_Qx_ = x 10^−4^) height scores. In comparison, scores that were computed using the UK Biobank (UKB) effect sizes showed substantially attenuated differences using both sub-sig (Q_x_ = 64, P_Qx_ = 5 x 10^−7^) and gw-sig (Q_x_ = 33, P_Qx_ = 0.02) SNPs, and a smaller difference between the two scores. This suggests that the attenuation of the signal in GIANT is not only driven by a loss of power when using fewer gw-sig SNPs, but also reflects a decrease in stratification effects. The overdispersion signal disappeared entirely when the UK Biobank family based effect sizes were used (**Figure 4, Supplementary Figure S16–S17**). Moreover, Q_x_ P values based on randomly ascertained SNPs and UK Biobank summary statistics are not uniformly distributed as would be expected if the theoretical null model is valid and if population structure is absent (**Supplementary Figure S19**). The possibility of residual stratification effects even in the UK Biobank is also supported by a recent study [23]. Therefore, we remain cautious about interpreting any residual signals as “real” signals of polygenic adaptation.

**Figure 4.**
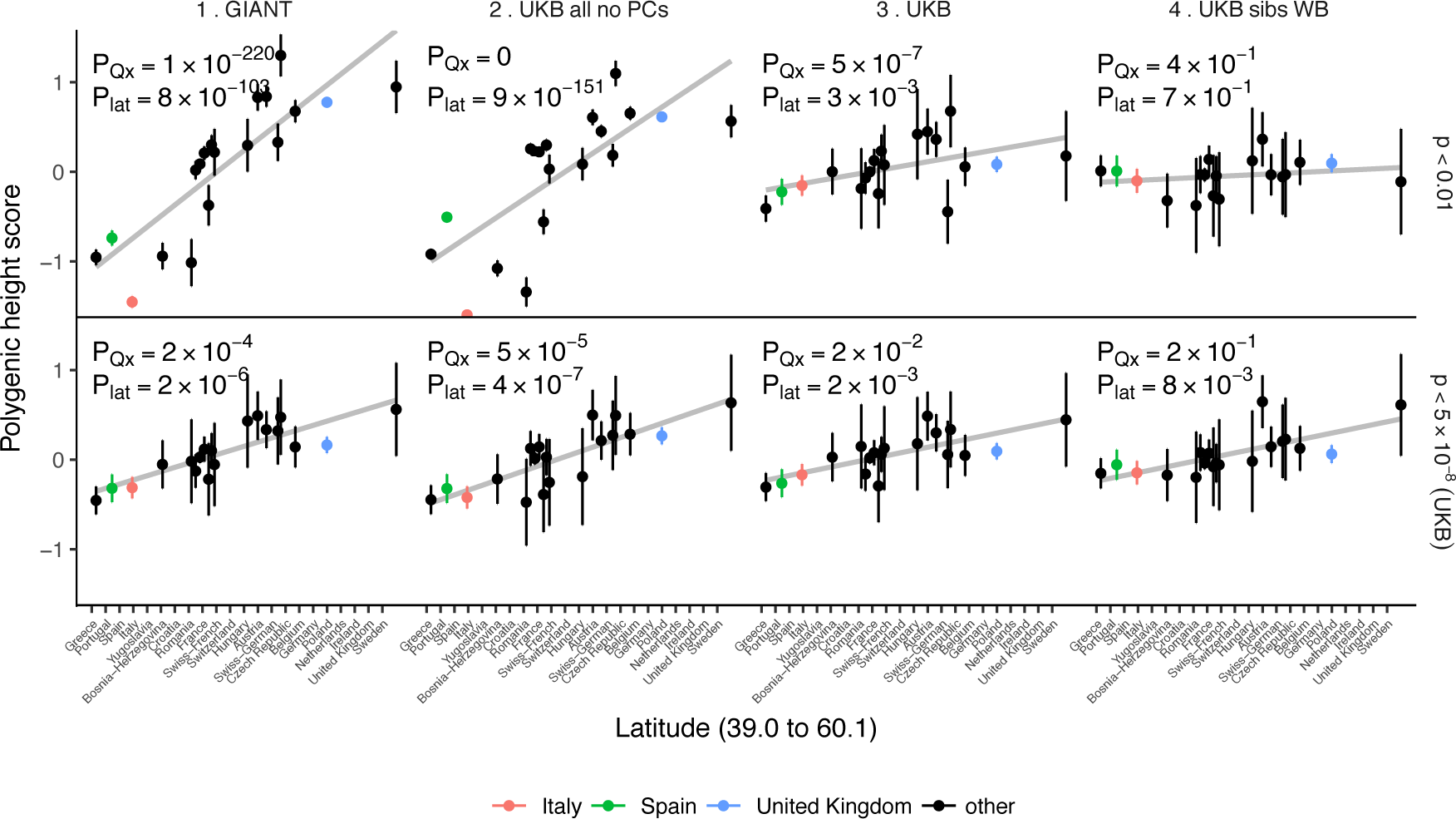
Polygenic height scores in POPRES populations show a residual albeit attenuated signal of polygenic adaptation for height. Standardized polygenic height scores from six summary statistics for 19 POPRES populations with at least 10 samples per population, ordered by latitude (see **Supplementary Table S3**). The grey line is the linear regression fit to the mean polygenic scores per population. Error bars represent 95% confidence intervals and are calculated in the same way as in **Figure 1**. SNPs which were overlapping between each set of the summary statistics and the POPRES SNPs were clumped using PLINK 1.9 with parameters r^^^2 < 0.1, 1 Mb distance, P < 1. (Top) A number of independent SNPs was chosen for each summary statistic to match the number of SNPs which remained when clumping UKB at P < 0.01. (Bottom) A set of independent SNPs with P < 5 x 10^−8^ in the UK Biobank was selected and used to compute polygenic scores along with effect size estimates from each of the different summary statistics. SNP numbers for each panel (P < 0.01 / P < 5 x 10^−8^): GIANT: 9463 / 869; UKB all no PCs: 9341 / 359; UKB: 9341 / 1034; UKB sibs WB: 9391 / 380. The numbers on each plot show the Q_x_ P value and the latitude covariance P value respectively for each summary statistic. See **Supplementary Figures S16-S18** for other clumping strategies and GWAS summary statistics.

## Discussion

We have shown that estimates of population differences in polygenic height scores are strikingly attenuated with the UK Biobank GWAS data relative to previous analyses. We find some evidence for population-level differences in genetic height, but it can only be robustly seen at highly significant SNPs, because any signal at less significant P values is dominated by the effect of residual population structure. Even genome-wide significant SNPs in these analyses may be subtly affected by population structure, leading to continued overestimation of the effect. Thus, it is difficult to arrive at any quantitative conclusion regarding the proportion of the population differences that are due to statistical biases vs. population stratification of genetic height. It is equally challenging to test whether differences in genetic height are due to adaptation in response to environmental differences, migration and admixture (e.g. fraction of Steppe pastoralist ancestry), or relaxation of negative selection. Further, estimates of the number of independent genetic loci contributing to height variation are sensitive to and likely confounded by residual population stratification.

We conclude that while effect estimates are highly concordant between GIANT and the UK Biobank when measured individually (**Supplementary Tables S4–S6, Supplementary Figure S1**), they are also influenced by residual population stratification that can mislead inferences about polygenic selection across populations in aggregate. Although these biases are subtle, in the context of tests for polygenic adaptation, which are driven by small systematic shifts in allele frequency, they can create highly significant artificial signals especially when SNPs that are not genome-wide significant are used to estimate genetic height. In no way do our results question the reliability of the genome-wide significant associations discovered in the GIANT cohort or the validity of the statistical methodology used in previously reported polygenic tests for adaptation. However, we urge caution in the interpretation of genome-wide signals of polygenic adaption that are based on large number of sub-significant SNPs-particularly when using effect sizes derived from meta-analysis of heterogeneous cohorts which may be unable to fully control for population structure.

## Materials and Methods

### Genome-wide association studies (GWAS)

We analyzed height using publicly available summary statistics that were obtained either by meta-analysis of multiple GWAS or by a GWAS performed on a single large population. We used results from the GIANT Consortium (N=253,288)[14] and a GWAS performed on individuals of the UK Biobank (UKB Neale” or simply “UK Biobank (UKB)”, N=336,474)[15] who derive their ancestry almost entirely from the British Isles (identified as “white British ancestry” by the UK Biobank). We also used an independent GWAS that included all UK Biobank individuals irrespective of ancestry and relatedness (“UKB Loh”, N=459,327)[24]. The Neale lab’s GWAS uses a linear model with sex and 10 principal components as covariates. Loh *et al.’s* GWAS uses a BOLT-LMM Bayesian mixed model. Association signals from the three studies are generally correlated for SNPs that are genome-wide significant in GIANT (see [[25]]).

We also used previously published family-based effect size estimates[7] (“NG2015 sibs”) as well as a number of test summary statistics on the UK Biobank that we generated to study the effects of population stratification. These are: “UKB Neale new” (Similar to UKB Neale, with less stringent ancestry definition and 20 PCs calculated within sample), “UKB all no PCs” (All UK Biobank samples included in the GWAS without correction by principal components), “UKB all 10 PCs” (All UK Biobank samples included in the GWAS with correction by 10 principal components), “UK WB no PCs” (Only “white British ancestry” samples included in the GWAS without correction by principal components), “UKB WB 10 PCs” (Only “white British ancestry” samples included in the GWAS with correction by 10 principal components), “UKB sibs all” (All UK Biobank siblings included in the GWAS), “UKB sibs WB” (Only UK Biobank “white British ancestry” siblings included in the GWAS) (Please see **Supplementary Table S1** for sample sizes and other details).

### Population genetic data for ancient and modern samples

We analyzed ancient and modern populations for which genotype data are publicly available. For ancient samples[26,27], we computed scores after dividing populations into three previously described broad ancestry labels (HG = Hunter-Gatherer (n=162 individuals), EF = Early Farmer (n=485 individuals), and SP = Steppe Ancestry (n=465 individuals)). For modern samples available through the 1000 genomes phase 3 release[28], we computed scores in 2 populations each from Northern Europe (GBR, CEU), Southern Europe (IBS, TSI), Africa (YRI, LWK), South Asia (PJL, BEB) and East Asia (CHB, JPT) (**Figure 1a, Supplementary Figures S2–S6**). In total, we analyzed 1112 ancient individuals, and 1005 modern individuals from 10 different populations in the 1000 genomes project (**Supplementary Table S2**). We used the allele frequency differences between the GBR and TSI populations for a number of analyses to study population stratification (**Figure. 3c, 3d, Supplementary Figures S11–S14**). We also analyzed 19 European populations from the POPRES[22] dataset with at least 10 samples per population (**Figure 4, Supplementary Table S3, Supplementary Figures S16–S18**).

All ancient samples had ‘pseudo-haploid’ genotype calls at 1240k sites generated by selecting a single sequence randomly for each individual at each SNP^9^. Thus, there is only a single allele from each individual at each site, but adjacent alleles might come from either of the two haplotypes of the individual. We also recomputed scores in present-day 1000 genomes individuals using only pseudohaploid calls at 1240k sites to allow for a fair comparison between ancient and modern samples (**Supplementary Figure S6**).

### Polygenic scores

The polygenic scores, confidence intervals and test statistics (against the null model of genetic drift) were computed based on the methodology developed in refs ([5],[29]). We computed the polygenic score (Z) for a trait in a population by taking the sum of allele frequencies across L sites associated with the trait, weighting each allele’s frequency by its effect on the trait (β_*l*_).

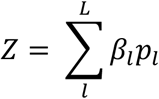

Al polygenic scores are plotted in centered standardized form 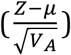, where 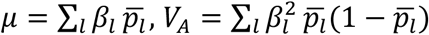, and 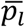 is the mean allele frequency across all populations analyzed.

Polygenic scores were computed using independent GWAS SNPs associated with height in three main ways: (1) The genome was divided into ~1700 nonoverlapping linkage disequilibrium (LD) blocks[30], and the SNP with the lowest P value within each block was picked to give a set of ~1700 independent SNPs for each height GWAS used (all SNPs for which effect sizes are available were considered) similarly to ref. [29]. In (2) and (3), Plink’s[31,32] clumping procedure was used to make independent “clumps” of SNPs for each GWAS at different P value thresholds. This procedure selects SNPs below a given P value threshold as index SNPs to start clumps around, and then reduces all SNPs (P < 0.01) that are in LD with these index SNPs (above an r^2^ threshold, 0.1) and within a physical distance of them (1 Mb) into clumps with them. Clumps are preferentially formed around index SNPs with the lowest P value in a greedy manner. The index SNP from each clump is then picked for further polygenic score analyses. The algorithm is also greedy such that each SNP will only appear in one clump if at all. We clumped each GWAS to obtain (2) a set of independent sub-significant SNPs associated with height (P < 0.01) similarly to ref. [7], and (3) a set of genome-wide significant SNPs associated with height (P < 5 x 10^−8^). The 1000 genomes phase 3 dataset was used as the reference panel for computing LD for the clumping procedure.

The estimated effect sizes for these three sets of SNPs from each GWAS was used to compute scores. Only autosomal SNPs were used for all analyses to avoid creating artificial mean differences between populations with different numbers of males and females.

The 95% credible intervals were constructed by assuming that the posterior of the underlying population allele frequency is independent across loci and populations and follows a beta distribution. We updated a Uniform prior distribution with allele counts from ancient and modern populations to obtain the posterior distribution at each locus in each population. We estimated the variance of the polygenic score *V*_*z*_ using the variance of the posterior distribution at each locus, and computed the width of 95% credible intervals as 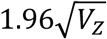 for each population.

The Q_x_ test statistic measures the degree of overdispersion of the mean population polygenic score compared to a null model of genetic drift. It assumes that the vector of mean centered mean population polygenic score follows a multivariate normal distribution: Z ~ MVN(0, 2 Va F), where Va is the additive genetic variance of the ancestral population and F is a square matrix describing the population structure. This is equivalent to the univariate case of the test statistic used in ref. [7]. The latitude test statistic assumes that Y’Z ~ N(0, 2 Va Y'FY), where Y is a mean centered vector of latitudes for each population.[33]

### tSDS analysis

The Singleton Density Score (SDS) method identifies signatures of recent positive selection based on a maximum likelihood estimate of the log-ratio of the mean tip-branch length of the derived vs. the ancestral allele at a given SNP. The tip-branch lengths are inferred from the average distance of each allele to the nearest singleton SNP across all individuals in a sequencing panel. When the sign of the SDS scores is aligned with the trait-increasing or trait-decreasing allele in the effect estimates of a GWAS, the Spearman correlation between the resulting tSDS scores and the GWAS P values has been proposed as an estimate of recent positive selection on polygenic traits.

Here, we applied the tSDS method using pre-computed Singleton Density Scores for 4,451,435 autosomal SNPs obtained from 3,195 individuals from the UK10K project [12,17] for SNPs associated with height in GIANT and the UK biobank (**Figure 1b**) and in different summary statistics (**Figure 3, Supplementary Figures S7–S8**). After normalizing SDS scores in each 1% allele frequency bin to mean zero and unit variance, excluding SNPs from the MHC region on chromosome 6 and aligning the sign of the SDS scores to the height increasing alleles (resulting in tSDS scores), we computed the Spearman correlation coefficient between the tSDS score and the GWAS P value. The tSDS Spearman correlation standard errors and P values were computed using a block-jackknife approach, where each block of 1% of all SNPs ordered by genomic location was left out and the Spearman correlation coefficient was computed on the remaining SNPs. We also compared the tSDS score distributions for only genome-wide significant SNPs (**Figure 3b**).

### Population structure analysis

To compute SNP loadings of the principal components of population structure (PC loadings) in the 1000 genomes data (**Figure 2, Supplementary Figure S10**), we first computed PC scores for each individual. We used SNPs that had matching alleles in 1000 genomes, GIANT and UK Biobank, that had minor allele frequency > 5% in 1000 genomes, and that were not located in the MHC locus, the chromosome 8 inversion region, or regions of long LD. After LD pruning to SNPs with r^2^ < 0.2 relative to each other, PCA was performed in PLINK on the 187,160 remaining SNPs. In order to get SNP PC loadings for more SNPs than those that were used to compute PC scores, we performed linear regressions of the PC scores on the genotype allele count of each SNP (after controlling for sex) and used the resulting regression coefficients as the SNP PC loading estimates.

## Acknowledgements

We thank Alkes Price, Jeremy Berg, Graham Coop, Jonathan Pritchard, Matthew Robinson, Jian Yang, Peter Visscher, Hilary Finucane, John Novembre and Raymond Walters for useful discussions and comments that significantly improved the manuscript. The study was supported by National Institute of Health grants HG009088, MH101244 (M.S., R.M., B.N. and S.S.) and GM127131 (S.S.). D.R. was supported by National Institutes of Health grant GM100233 and HG006399, an Allen Discovery Center of the Paul Allen Foundation, and the Howard Hughes Medical Institute.

## Supplementary Note

### Characterization of stratification effects in GIANT and UK Biobank

To better understand how stratification influences the differences observed between GIANT and UK Biobank, we grouped SNPs by their P value in GIANT and by their P value in UK Biobank (**Supplementary Figure S1**). First, we observe that SNPs with low GIANT P values, but not SNPs with low UK Biobank P values, show greater differences in estimated effect size (**Supplementary Figure S1a**). However, the relative difference in beta values decreases for lower P values, and the correlation among betas approaches one at the most significant SNPs (**Supplementary Figure S1b,c**). In GIANT, more significant SNPs exhibit a greater correlation between effect estimates and GBR-TSI allele frequency differences, while this is not observed in the UK Biobank (**Supplementary Figure S1d**). Consequently, the difference in UK Biobank and GIANT effect size estimates is more correlated to GBR-TSI allele frequency differences at more significant SNPs (**Supplementary Figure S1e**). This suggests that while stratification effects are larger at more significant SNPs, the magnitude of stratification-independent effects is even larger, which may be why polygenic score results converge when using only the most significant SNPs.

Next, we investigated how P value inflation as measured by *X*_*GC*_ is influenced by stratification, by grouping SNPs into deciles based on their GBR-TSI allele frequency difference (**Supplementary Figure S12**). To guard against the effect of observing lower P values at more differentiated SNPs simply because those SNPs are more common on average, we restrict this analysis to SNPs with mean MAF > 20%. We find that *X*_*GC*_ is not much increased for SNPs that are more differentiated between populations. However, in the presence of stratification, there is a large difference between *X*_*GC*_ of height increasing alleles and *X*_*GC*_ of height decreasing alleles (**Supplementary Figure S12b**). Similarly, there are large effects on the frequency with which a SNP is estimated to be height increasing or height decreasing. In GIANT, SNPs in the highest decile of GBR-TSI allele frequency differences are 52% more often estimated to be height increasing than height decreasing, while these rates are close to even in the UK Biobank (**Supplementary Figure S12a**).

### Supplementary figures

**Figure S1.**
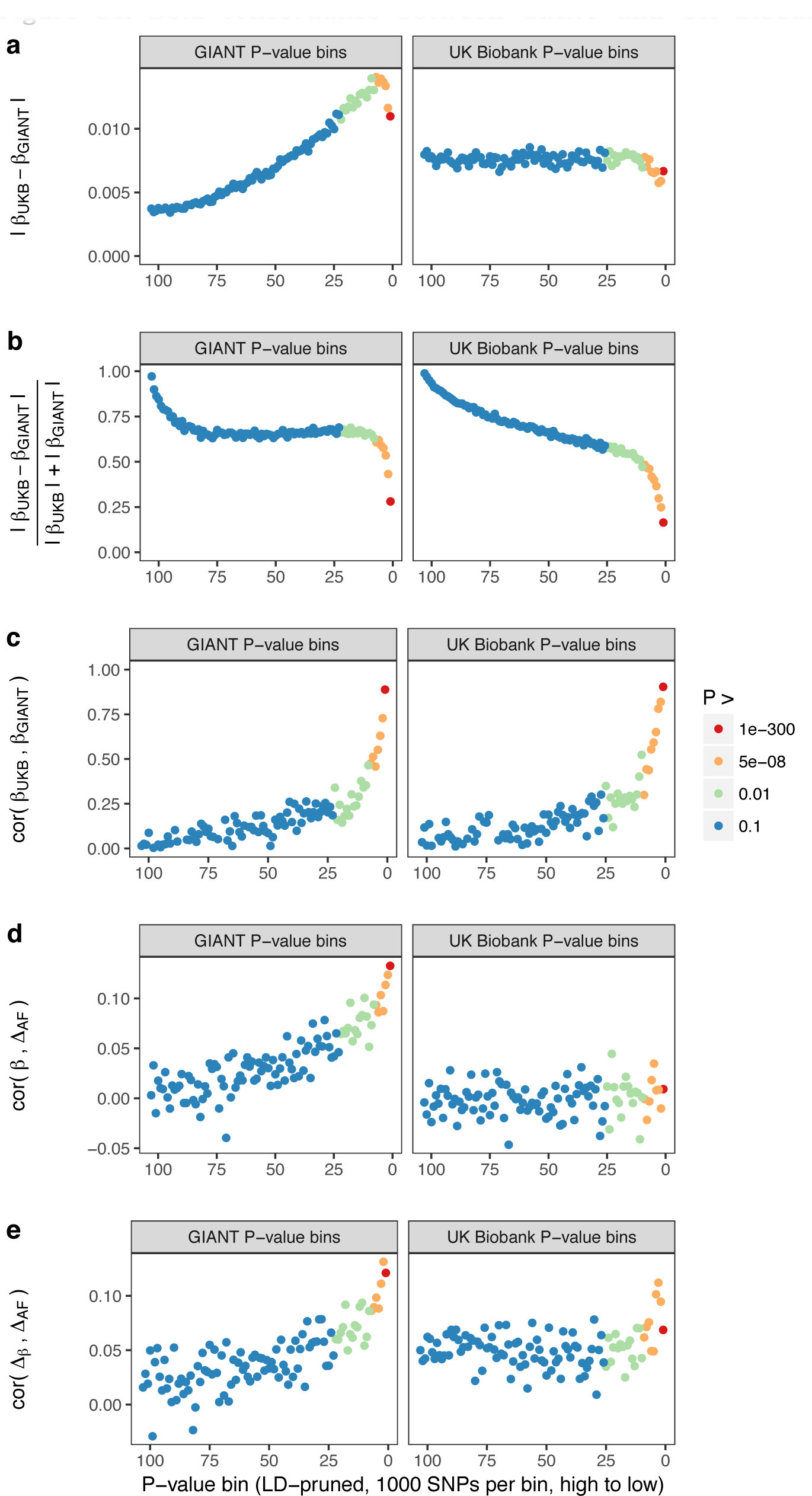
Beta concordance between GIANT and UK Biobank by P value bin. SNPs intersecting between GIANT and UKB were LD-pruned (using PLINK 1.9 with parameters r^^^2 = 0.1, window size = 1 Mb, step size 5) and grouped into P value bins of 500 SNPs each, for P values from GIANT (left) and UKB (right). Color is based on the smallest P value in each bin. (a) Absolute beta difference. As expected, absolute beta and thus the absolute beta difference increases across P value bins. (b) Absolute beta difference, scaled by the sum of absolute betas. The relative difference of absolute betas decreases for lower P values. (c) Pearson correlation among betas approaches one for the lowest P values. (d) Correlation between beta (left GIANT, right UK Biobank) and GBR-TSI allele frequency difference. (e) Correlation between the GIANT - UK Biobank beta difference and GBR-TSI allele frequency difference.

**Figure S2.**
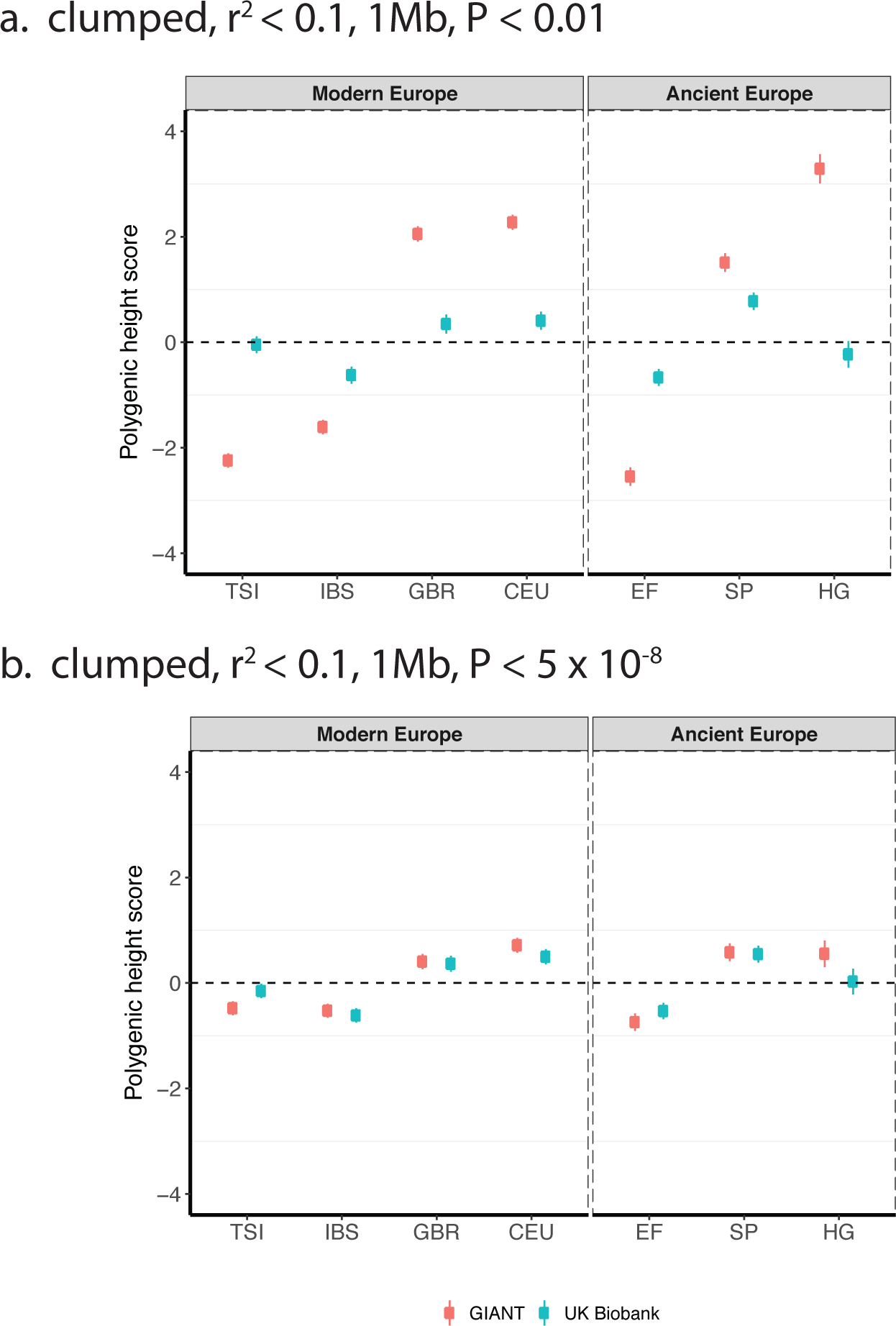
Polygenic scores using height-associated SNPs from GIANT- and UK Biobank-based GWA studies for clumped SNPs in present-day and ancient Europeans. Scores are shown, centered by the average score across modern and ancient populations respectively and standardized by the square root of the additive variance. SNPs were LD-pruned with plink's clumping procedure for parameters: (a] r^2^ < 0.1, 1Mb, P < 0.01 (81,941 SNPs in UKB, 22,561 SNPs in GIANT], and (b) r^2^ < 0.1, 1Mb, P < 5×10^−8^ (4478 SNPs in UKB, 1442 SNPs in GIANT). Modern populations are shown from Northern Europe (CEU, GBR] and Southern Europe (IBS, TSI] from the 1000 genomes project. Ancient populations are shown in three meta-populations, hunter-gatherers (HG), early farmers (EF) and Steppe Ancestry (SP). Error bars are drawn at 95% credible intervals.

**Figure S3.**
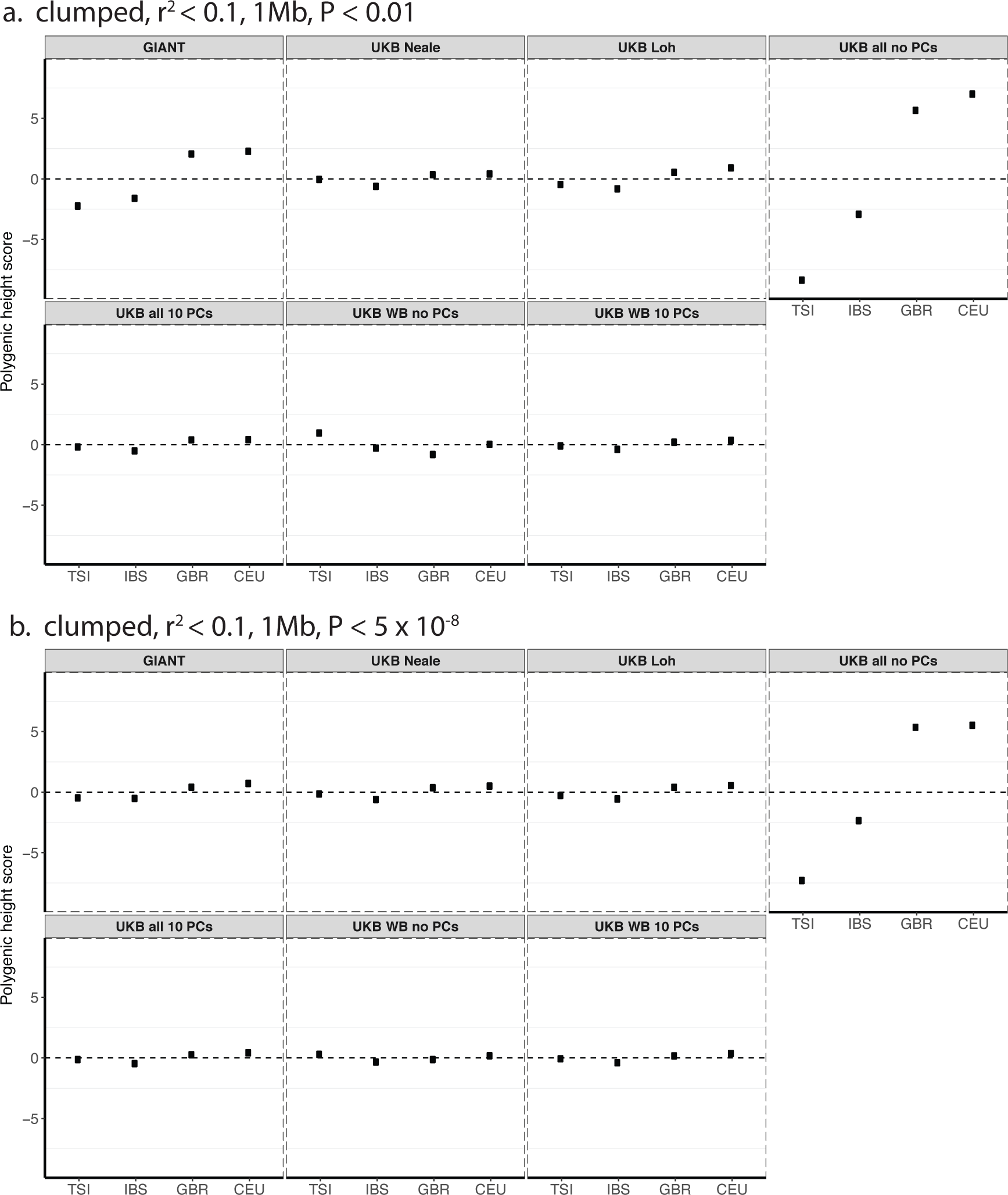
Polygenic height scores in 1000 genomes European populations using clumped SNPs and effect sizes from different summary statistics. Polygenic scores in modern European populations are shown using SNPs LD-pruned with PLINK’s clumping procedure with parameters: (a) r^2^ < 0.1, 1Mb, P < 0.01, and (b) r^2^ < 0.1, 1Mb, P < 5×10^−8^. Scores are centered by the average score across populations and standardized by the square root of the additive variance. Modern populations are shown from Northern Europe (CEU, GBR) and Southern Europe (IBS, TSI) from the 1000 Genomes Project. Error bars are drawn at 95% credible intervals.

**Figure S4.**
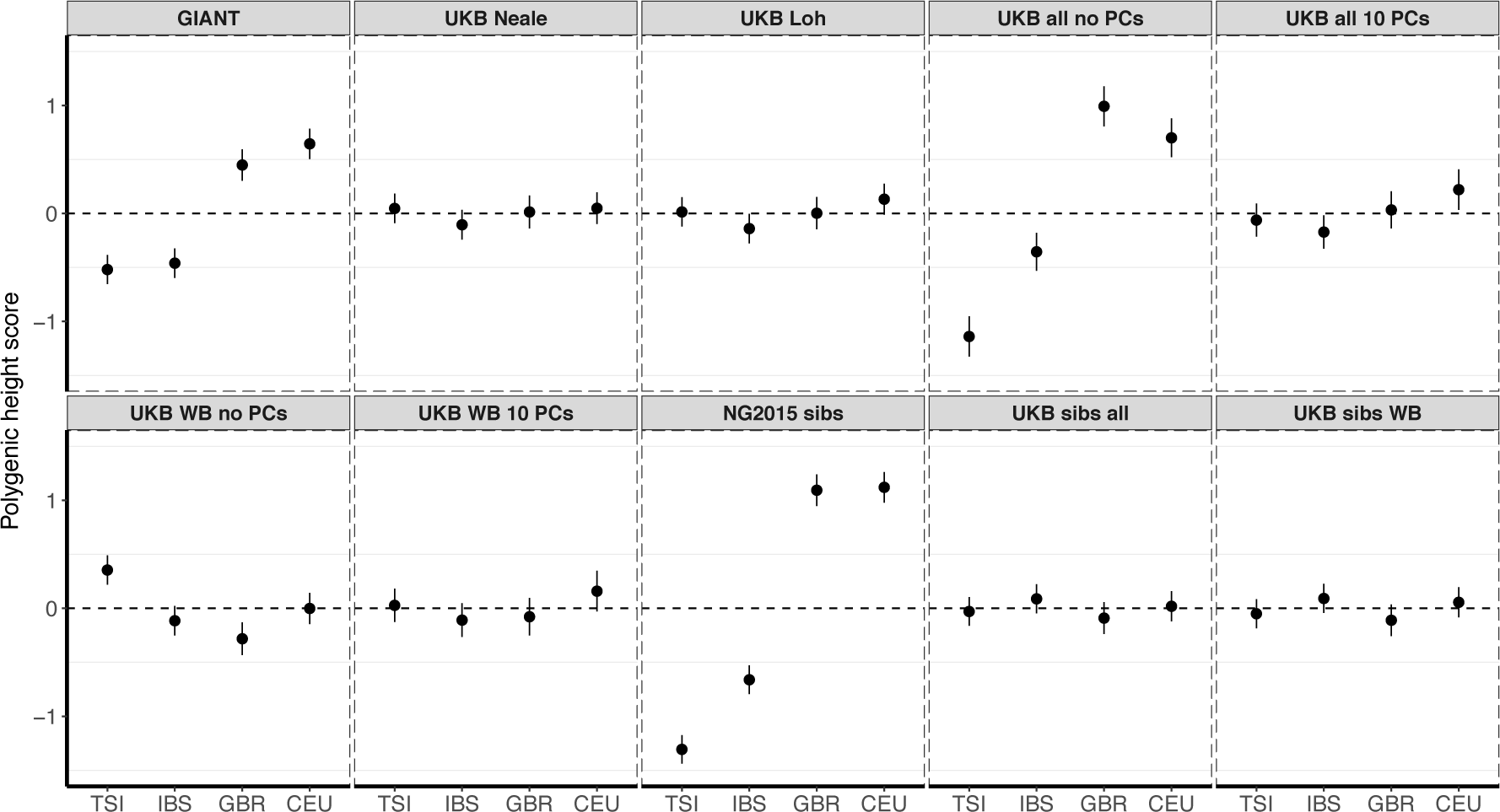
Polygenic height scores in 1000 Genomes Project European populations using ~1700 independent SNPs and effect sizes from different summary statistics. Polygenic scores in modern European populations are shown using SNPs LD-pruned by picking the SNP with the lowest P value in each of ~1700 LD-independent blocks genome-wide. Scores are centered by the average score across populations and standardized by the square root of the additive variance. Modern populations are shown from Northern Europe (CEU, GBR] and Southern Europe (IBS, TSI] from the 1000 Genomes Project. Error bars are drawn at 95% credible intervals.

**Figure S5.**
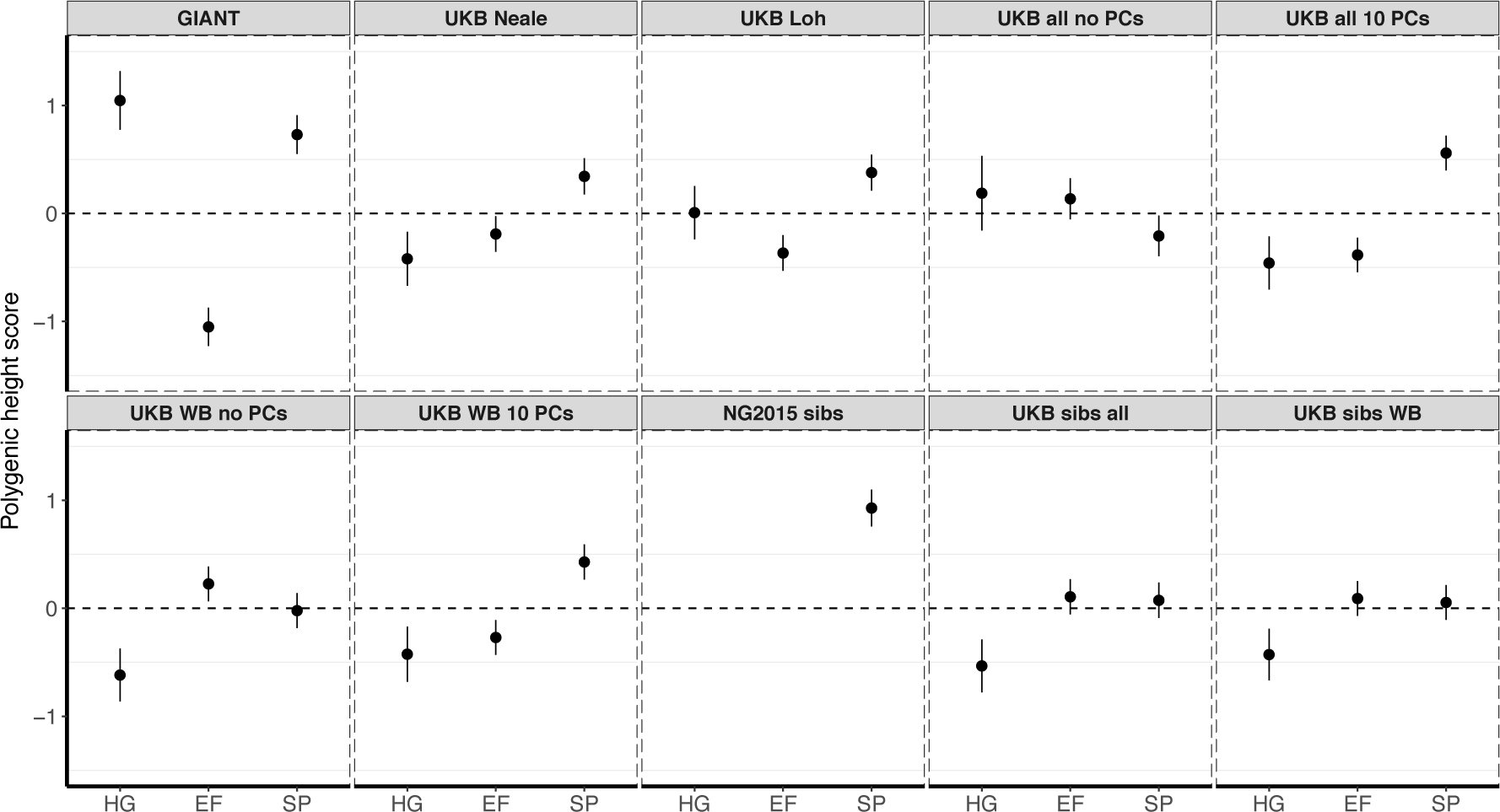
Polygenic height scores in ancient populations using ~1700 independent SNPs and effect sizes from different summary statistics. Polygenic scores in ancient meta-populations are shown using SNPs LD-pruned by picking the SNP with the lowest P value in each of ~1700 LD-independent blocks genome-wide. Scores are centered by the average score across populations and standardized by the square root of the additive variance. Error bars are drawn at 95% credible intervals̤ Ancient populations are shown in three meta-populations (HG = Hunter-Gatherer (n=162 individuals), EF = Early Farmer (n=485 individuals), and SP = Steppe Ancestry (n=465 individuals)). The y-axis is truncated at (−1.5, 1.5) for all panels - this omits two points in the NG2015 sibs panel: HG [3.86 (CI: 3.60, 4.12)], EF [−2.18(CI: −2.34, −2.02)].

**Figure S6.**
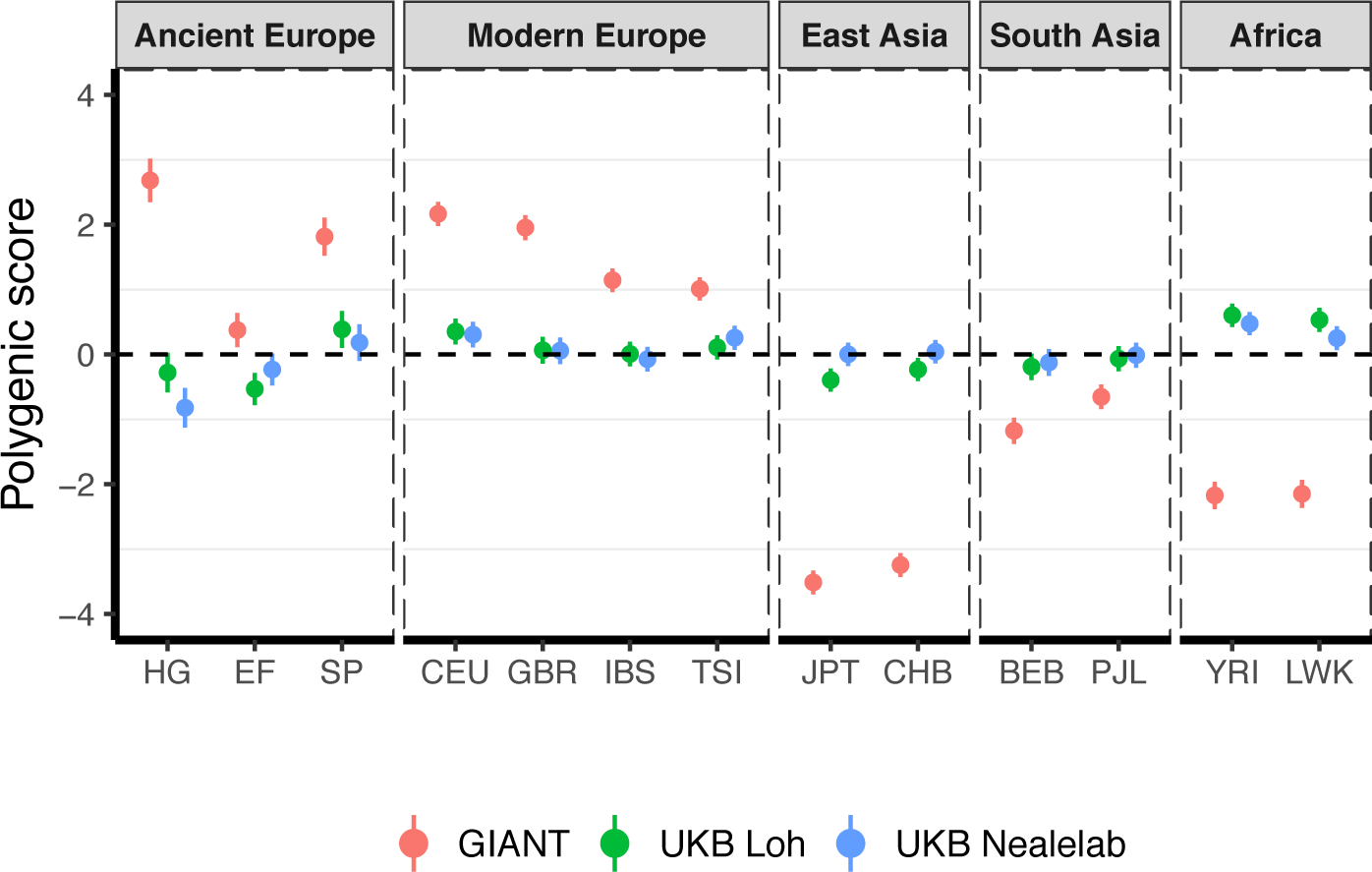
Polygenic height scores in ancient and global modern populations using three different GWAS. All scores are centered by the average score across all populations (μ_*GIANT*_= 0.645, μ_*LOH*_ = −0.219, μ_*NEALELAB*_= −0.259) and standardized by the square root of the additive variance. Error bars are drawn at 95% credible intervals. Modern populations are shown from Northern Europe (CEU, GBR), Southern Europe (IBS, TSI), South Asia (PJL, BEB), East Asia (CHB, JPT) and Africa (YRI, LWK). Ancient populations are shown in three metapopulations, hunter-gatherers (HG), early farmers (EF) and Steppe Ancestry (SP). Pseudohaploid genotype calls were made for modern populations before computing polygenic scores to allow fair comparison with ancient DNA. SNPs were LD-pruned by picking the SNP with the lowest P value in each of ~1700 LD-independent blocks genome-wide.

**Figure S7.**
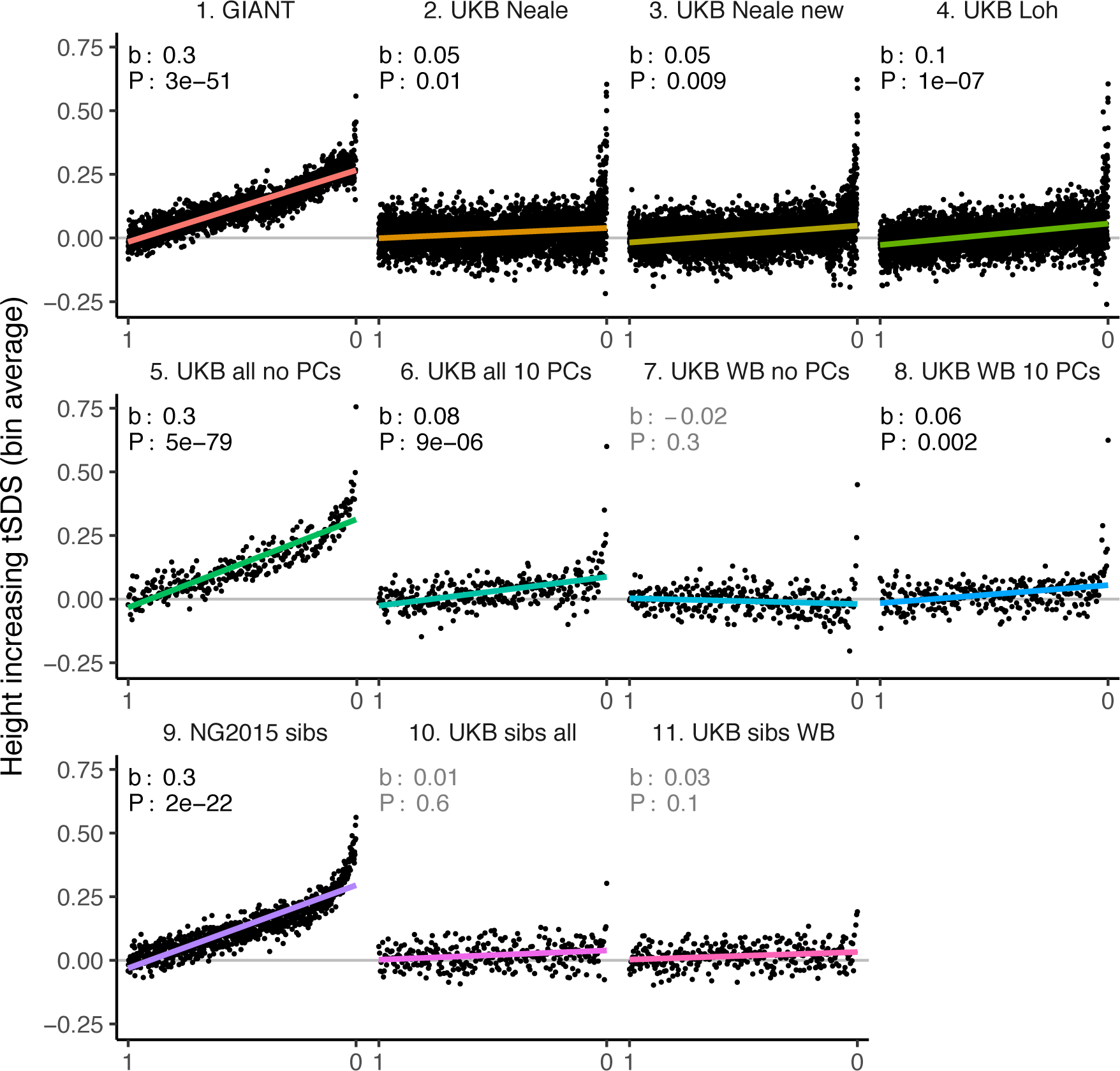
tSDS for height-increasing alleles using effect sizes from different summary statistics. SNPs were ordered by GWAS P value and grouped into bins of 1000 SNPs each. The mean tSDS score within each P value bin is shown on the y-axis. In contrast to Figure 3, where Spearman correlation coefficients and Jackknife standard errors were computed, here we show the regression slope and P value, which were computed on the un-binned data. The gray line indicates the null-expectation, and the colored lines are the linear regression fit. The lowest P value bin in panel 5 with a y-axis value of 1.5 has been omitted.

**Figure S8.**
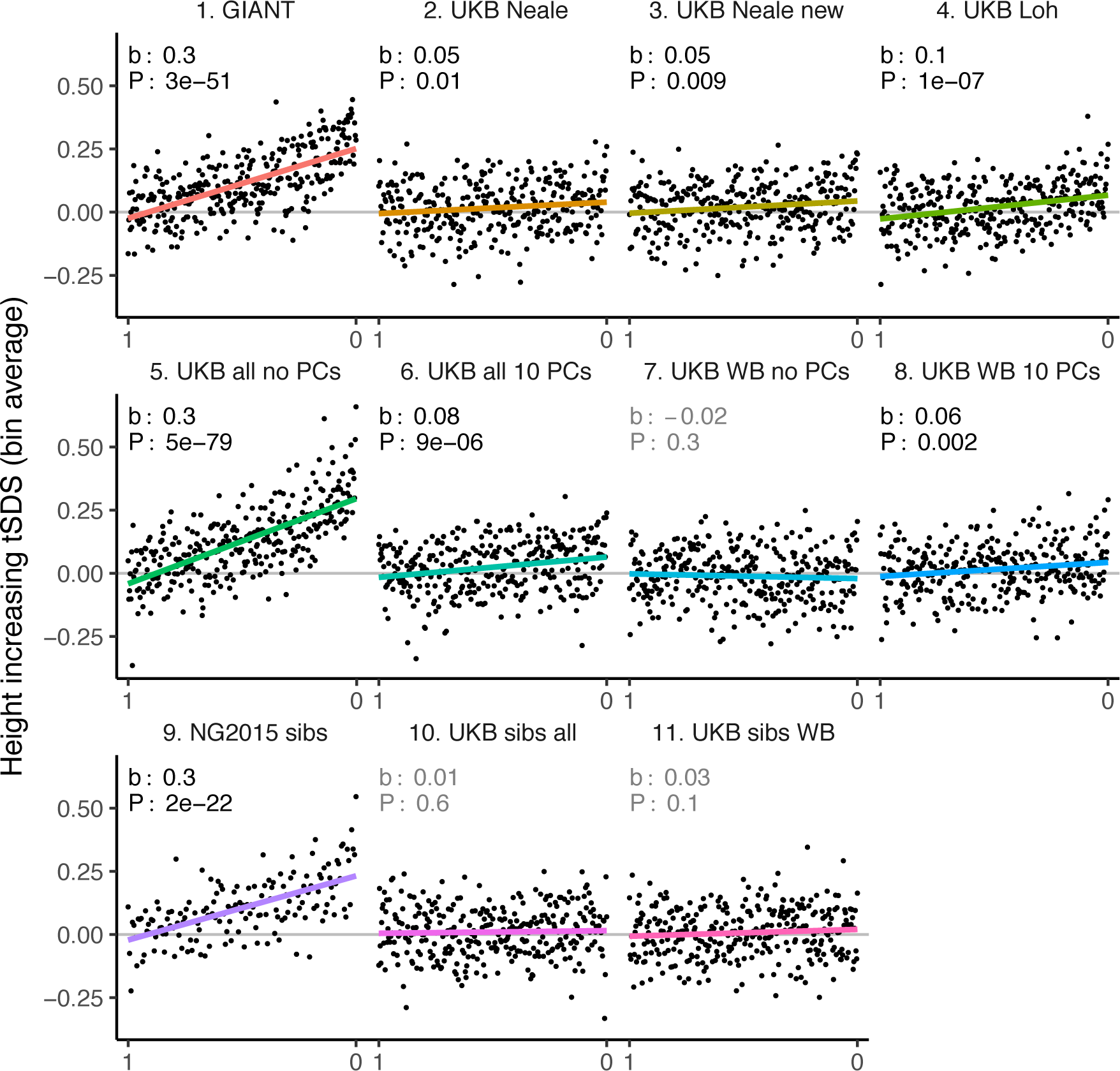
tSDS for LD-pruned height-increasing alleles using effect sizes from different summary statistics. Binning SNPs by P value can lead to spurious results at the low P value bins when SNPs are in LD (**Figure S15**) Here, LD-pruned SNPs were ordered by GWAS P value and grouped into bins of 100 SNPs each. The mean tSDS score within each P value bin is shown on the y-axis. In contrast to **Figure 3**, where Spearman correlation coefficients and Jackknife standard errors were computed, here we show the regression slope and P value, which were computed on the un-binned data. The gray line indicates the null-expectation, and the colored lines are the linear regression fit.

**Figure S9.**
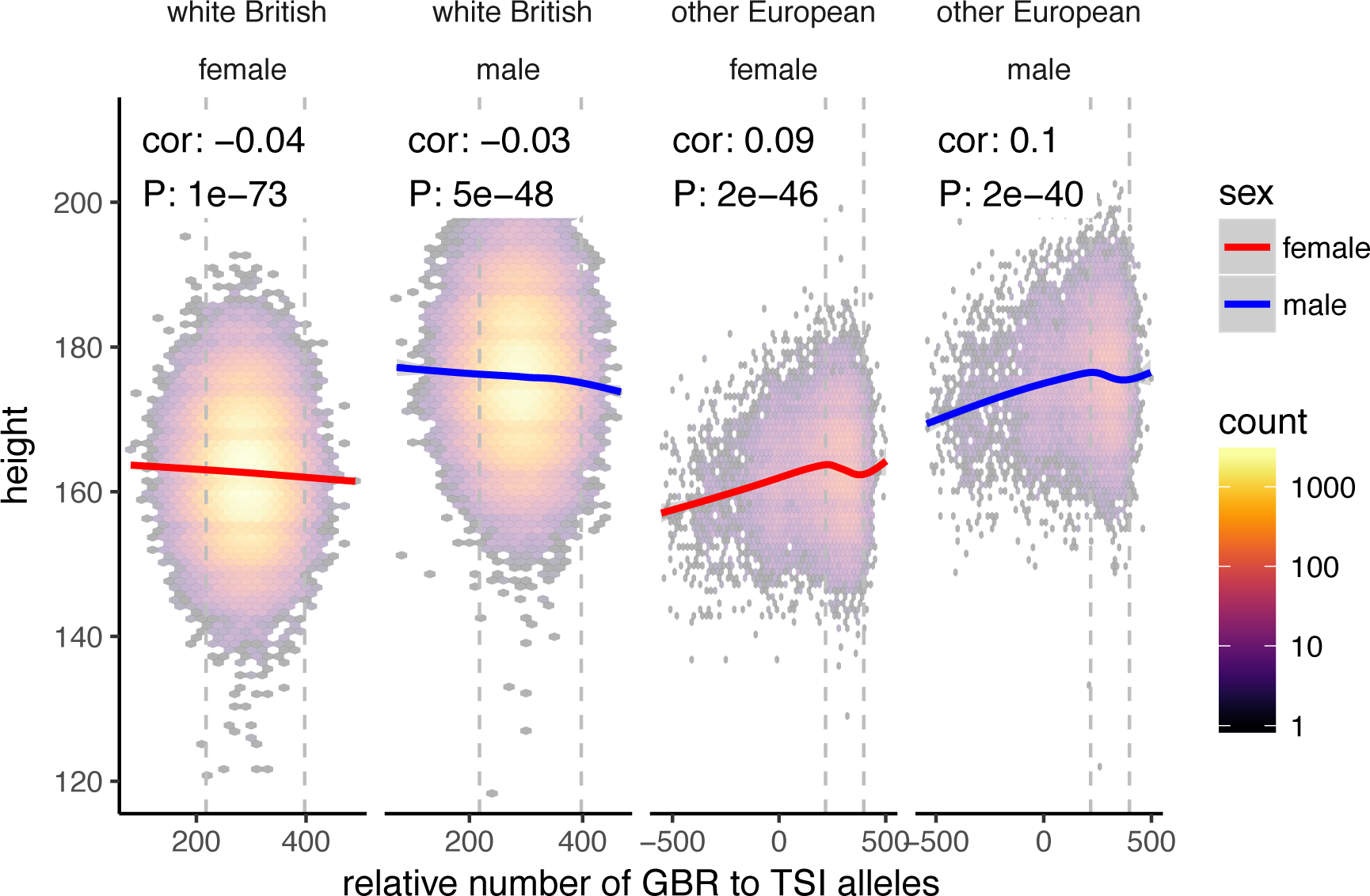
Height (cm) in the UKB as a function of GBR-TSI score. We computed the relative number of GBR to TSI related alleles in each sample by multiplying the allele frequency difference by the number of alternative alleles in each sample in the UKB (GBR-TSI score). Vertical lines indicate 5th and 95th percentile of among-white British samples, showing that there is a significant negative relationship between the GBR-TSI allele sharing score and height (in cm). Among all other broadly European samples, this relationship is significantly positive across the whole range, but again significantly negative in the white British range. This can explain why stratification effects go in opposite directions in a UKB height GWAS of white British samples and a UKB height GWAS of all samples. Here, other European samples were defined as those that lie within the mean +/− 24 standard deviations along the first six principal components.

**Figure S10.**
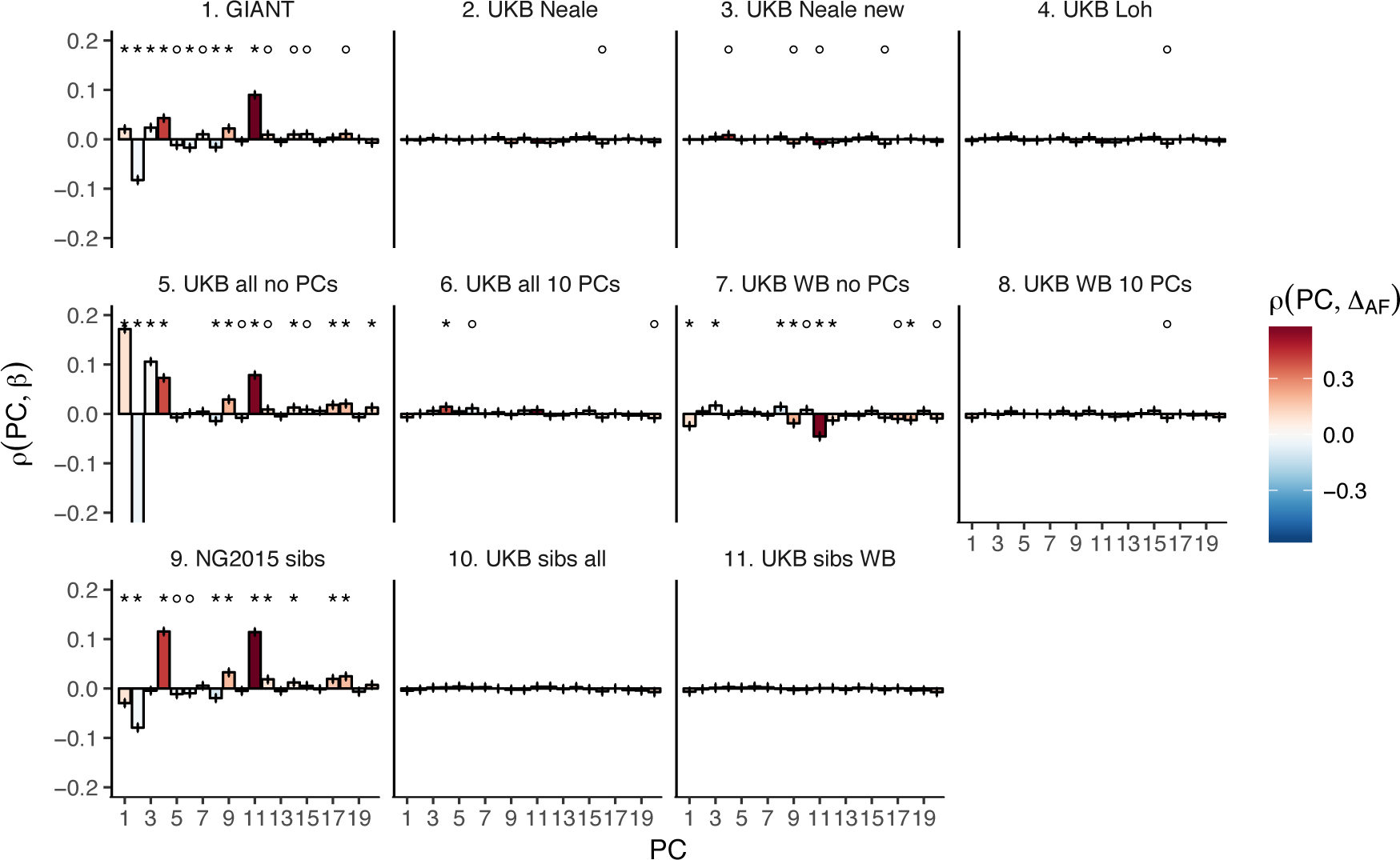
Pearson Correlation coefficients of PC loadings and height beta coefficients for different summary statistics. PCs were computed in all 1000 genomes phase 1 samples. Colors indicate the correlation of each PC loading with the allele frequency difference between GBR and TSI, a proxy for the European North-South genetic differentiation. PC 4 and 11 are most highly correlated with the GBR - TSI allele frequency difference. Error bars indicate 95% confidence interval of the correlation coefficient, assuming 60,000 independent genetic markers. We confirmed that the resulting errors are similar to block jackknife standard errors. Open circles indicate correlations significant at alpha = 0.05, stars indicate correlations significant after Bonferroni correction in 20 PCs (P < 0.0025).

**Figure S11.**
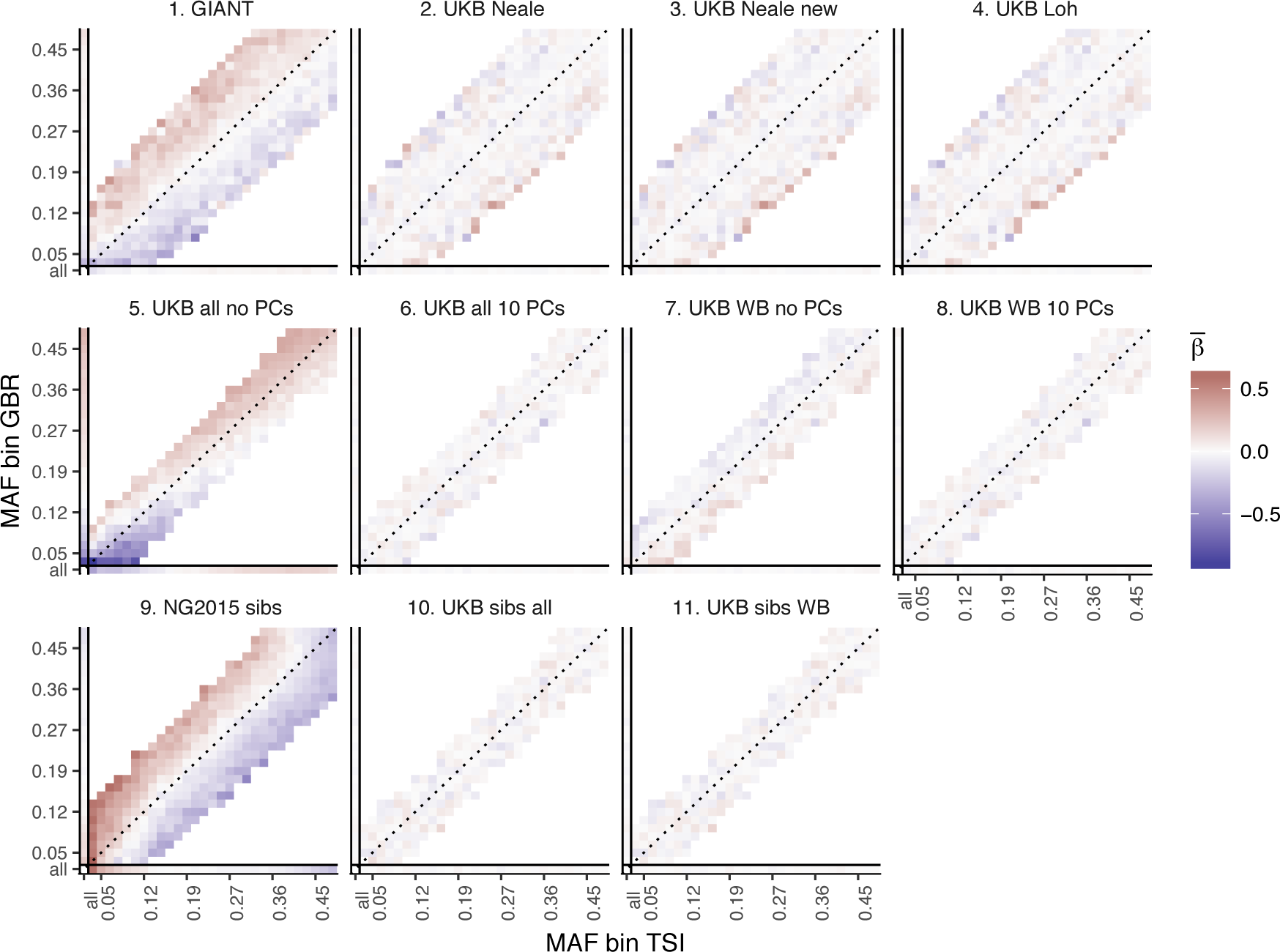
Heat map of mean beta coefficients for different summary statistics. All SNPs are binned by GBR and TSI minor allele frequency. Only bins with at least 300 SNPs are shown. Panel 7 (as well as 2, 3 and 4] shows stratification effects in opposite direction to those in GIANT. **Figure S9** illustrates how these opposite-direction stratification effects can arise.

**Figure S12.**
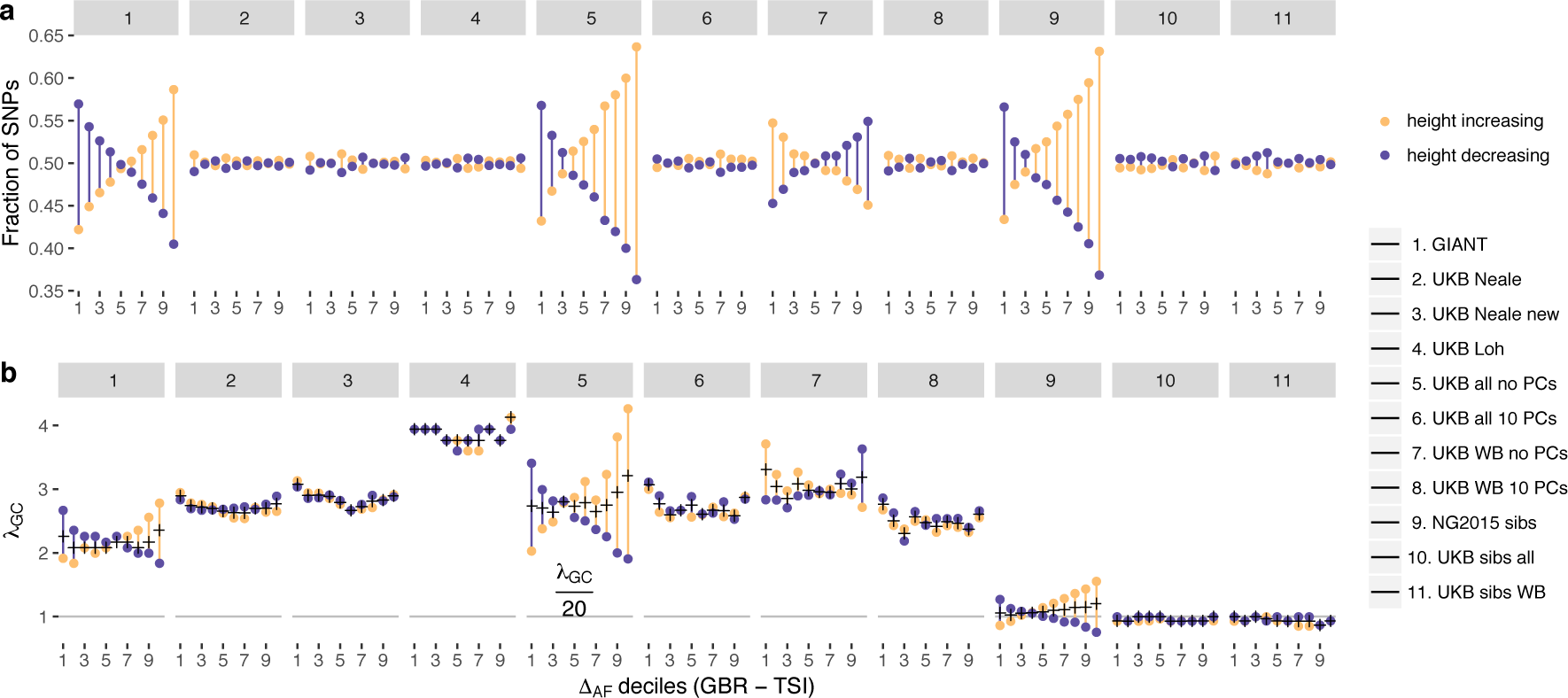
Effect of GBR-TSI allele frequency difference on beta estimates and P values. SNPs with MAF > 0.2 (based on mean between TSI and GBR) were grouped into GBR-TSI allele frequency difference deciles, with the first decile representing SNPs less common in GBR and the last decile representing SNPs more common in GBR. Fraction of height-increasing (yellow dots) vs. height-decreasing SNPs (purple dots) in each decile. In GIANT, 59% of SNPs in the highest decile are estimated to be height-increasing, and 41% are estimated to be height-decreasing. In the UK Biobank, this ratio is close to 50-50. Lambda-GC in each decile for height-increasing (yellow dots) vs. height-decreasing SNPs (purple dots). In GIANT, the median P value of SNPs in the highest decile is 2.78 for SNPs estimated to be height-increasing and 1.83 for SNPs estimated to be height-decreasing (a difference of 52%). In the UK Biobank, the median P value of SNPs in the highest decile is 2.65 for SNPs estimated to be height-increasing and 2.89 for SNPs estimated to be height-decreasing (a difference of only 9%, going in the opposite direction).

**Figure S13.**
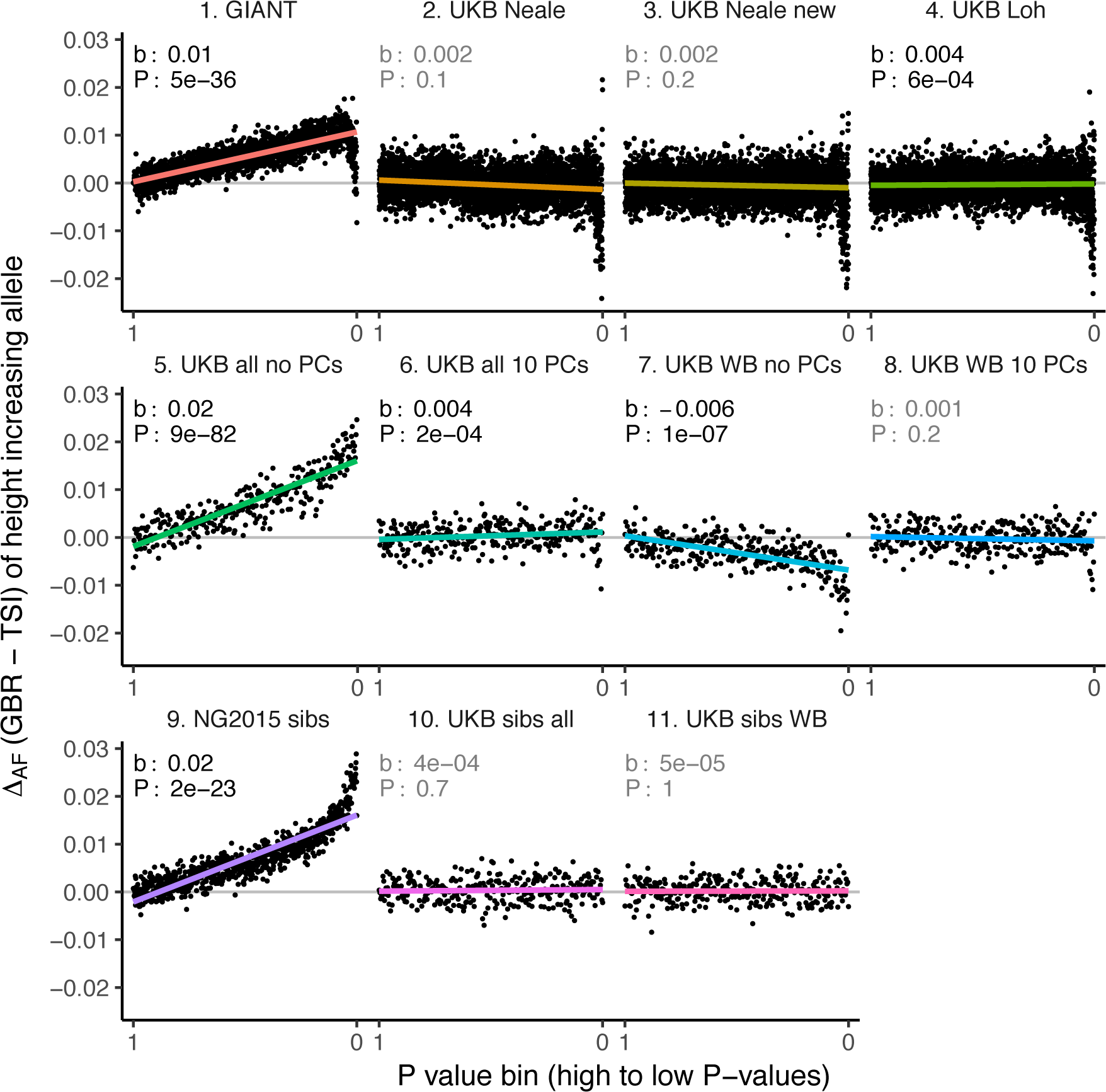
Allele frequency difference for height-increasing alleles using different summary statistics. SNPs were ordered by GWAS P value and grouped into bins of 1000 SNPs each. The gray line indicates the null-expectation, and the colored lines are the linear regression fit. The lowest P value bin in panel 5 with a y-axis value of 0.06 has been omitted.

**Figure S14.**
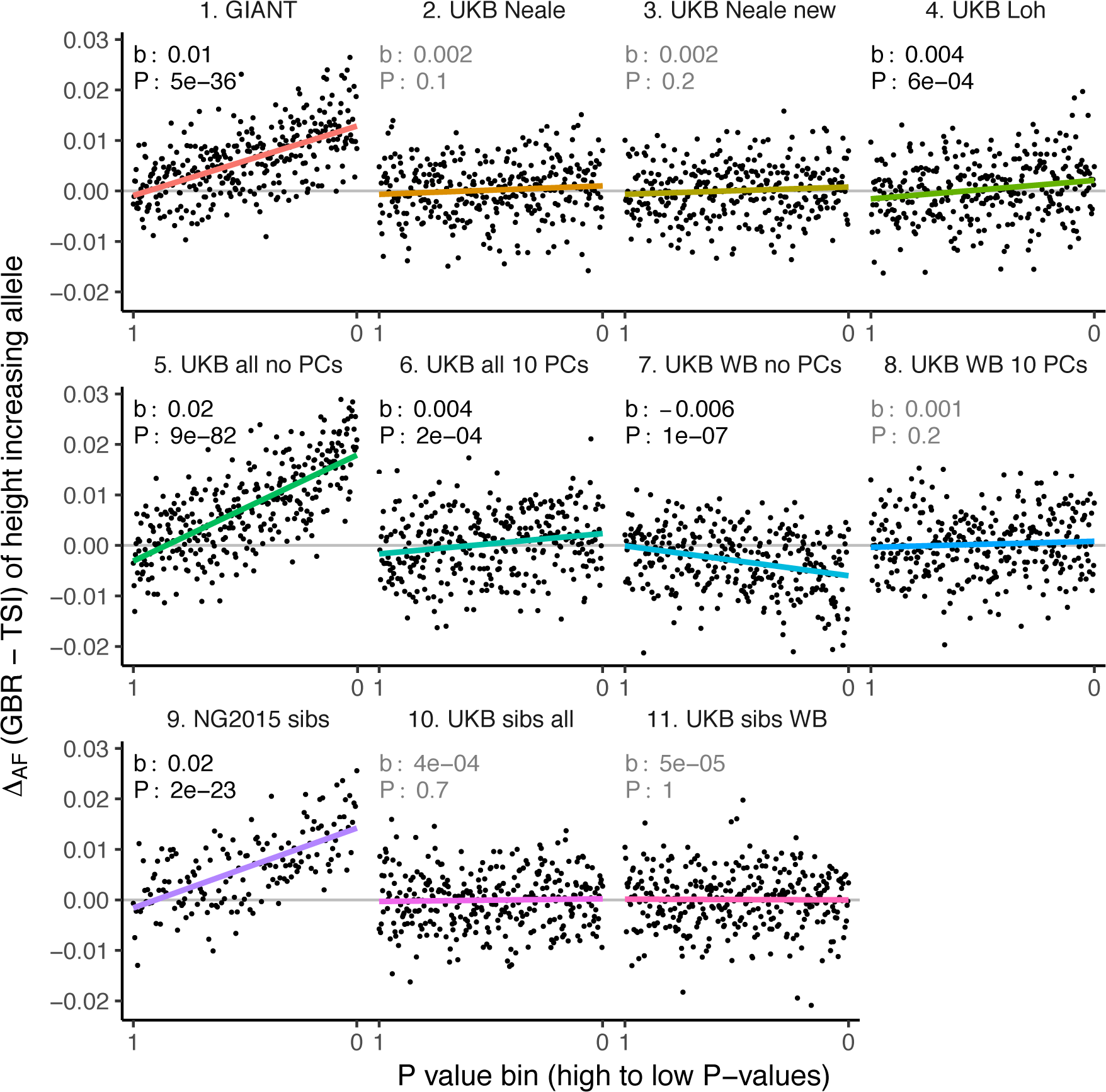
Allele frequency difference for LD-pruned height-increasing alleles using different summary statistics. Binning SNPs by P value can lead to spurious results at the low P value bins when SNPs are in LD (**Figure S15**) Here, LD-pruned SNPs were ordered by GWAS P value and grouped into bins of 100 SNPs each. The gray line indicates the null-expectation, and the colored lines are the linear regression fit.

**Figure S15.**
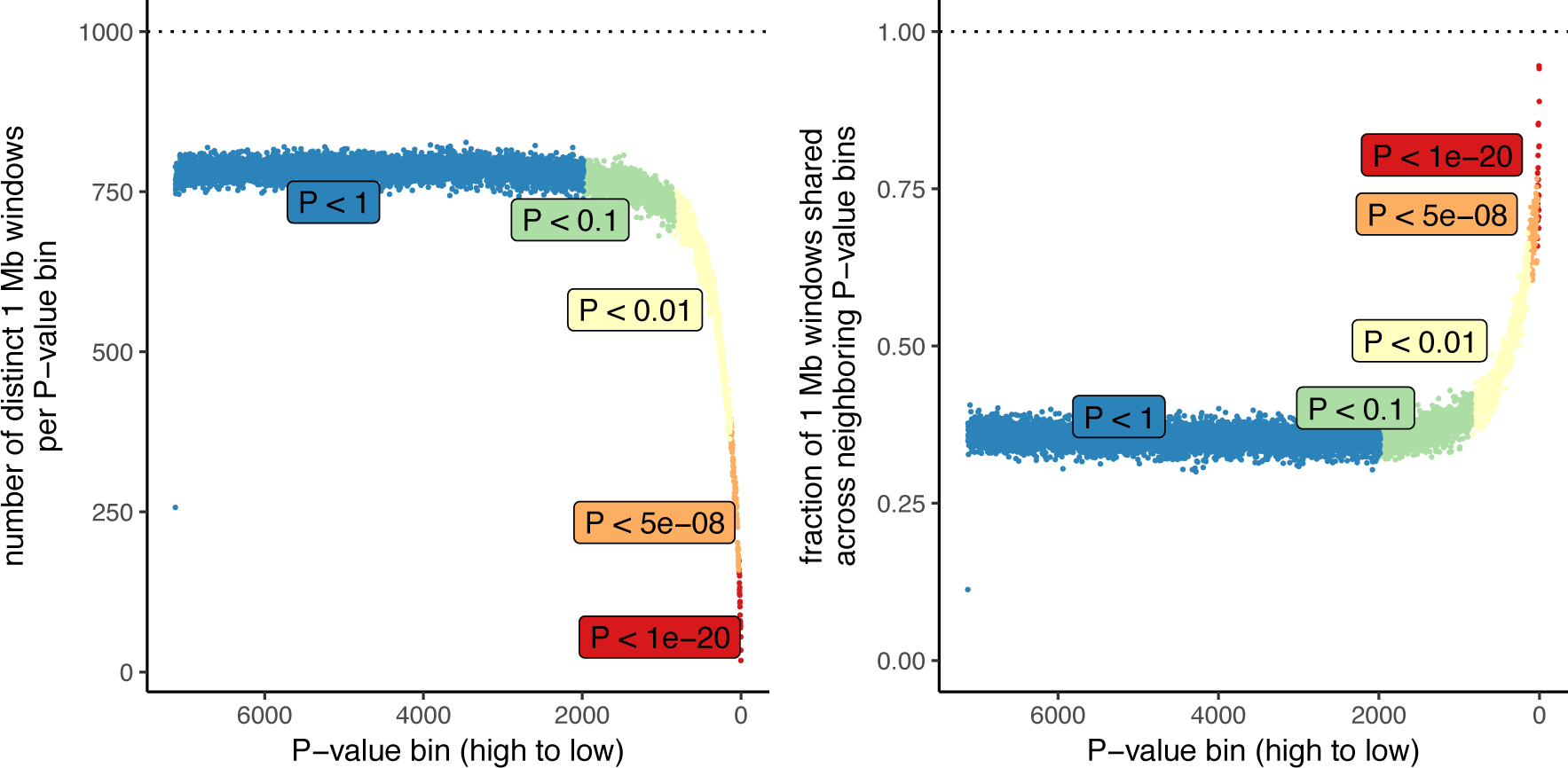
Number of independent regions per GWAS P value bin in the UK Biobank. SDS results in Field et al. as well as in **Figure 3** in this article are visualized by grouping non-independent SNPs into bins according to their P value. This may lead to unpredictable patterns at the low end of the P value distribution, because the lowest P value bins do not represent independent signals. This is demonstrated here, by grouping all UKB SNPs into bins of 1000 SNPs each, as in the SDS plots in **Figure 1b** and **Figure 3**. Left: The number of independent SNPs per P value bin is much lower at lower P values. Right: Neighboring P value bins share a large fraction of 1Mb regions at lower P values. This demonstrates that the lowest P value bins do not represent independent signals if SNPs are not LD-pruned and can exhibit patterns that are dominated by one or a few LD-regions.

**Figure S16.**
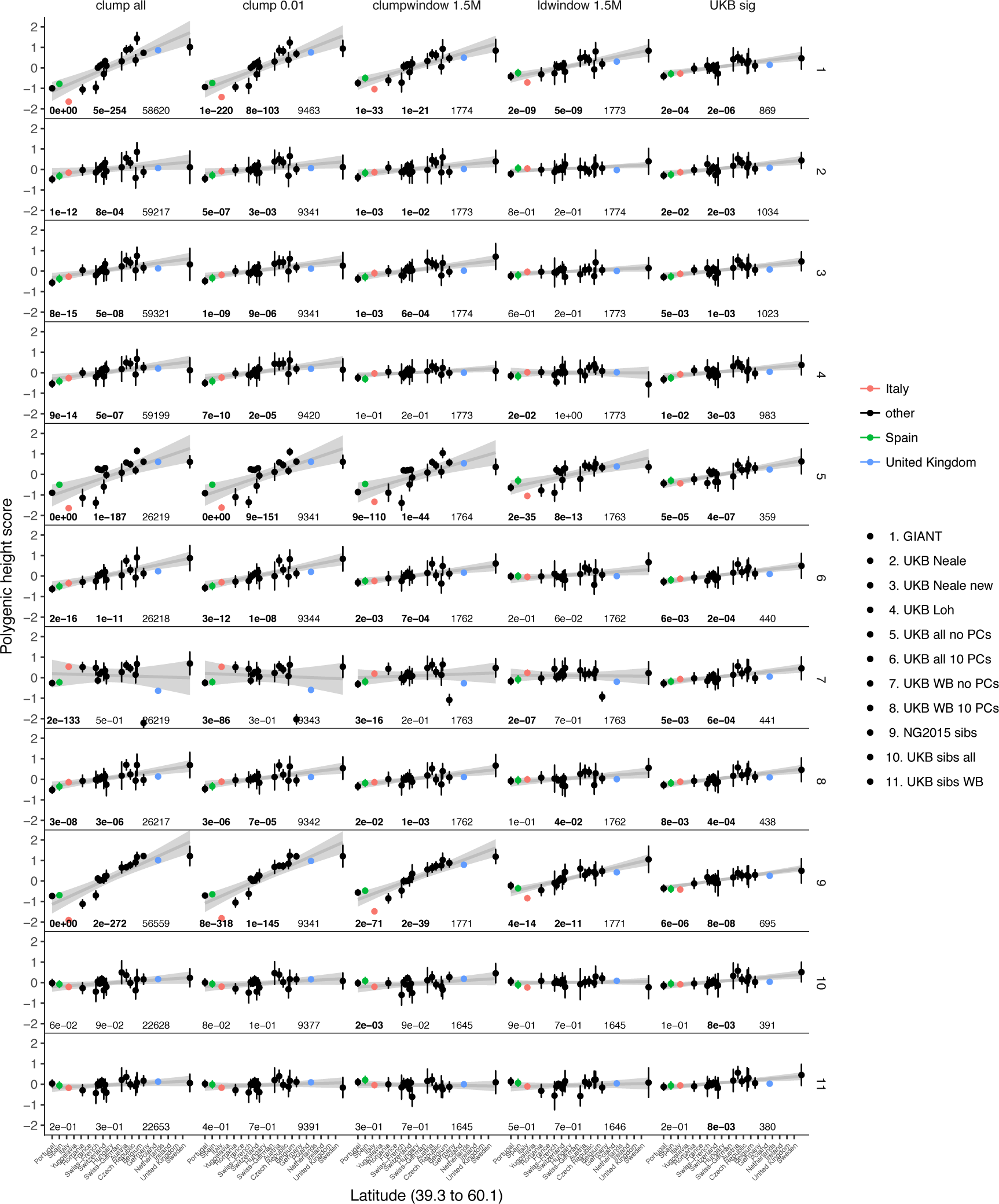
Polygenic height scores in POPRES for different summary statistics. Standardized polygenic height score from diverse summary statistics for 19 POPRES populations with at least 10 samples per population, ordered by latitude (see **Table S3**). Confidence intervals and clumping procedure are the same as in (a). The gray line is the linear regression fit to the mean polygenic height score per population. The numbers on each plot show the Q_x_ P value, the latitude covariance P value and the number of SNPs respectively for each summary statistic. Each column shows a different selection of SNPs. clump all: clumped SNPs with no P value threshold; clump 0.01: clumped SNPs with P < 0.01 in UKB and the same number of SNPs in other summary statistics (same as **Figure 4**); clumpwindow 1.5M: genome was split into blocks of 1.5 Mb, lowest P-value SNP was picked in each bin, similar to the 1700 blocks; ldwindow 1.5Mb: genome was split into blocks of 1.5 Mb, random SNP was picked in each bin; UKB sig: LD-pruned SNPs with P < 5×10^−8^ in UKB.

**Figure S17.**
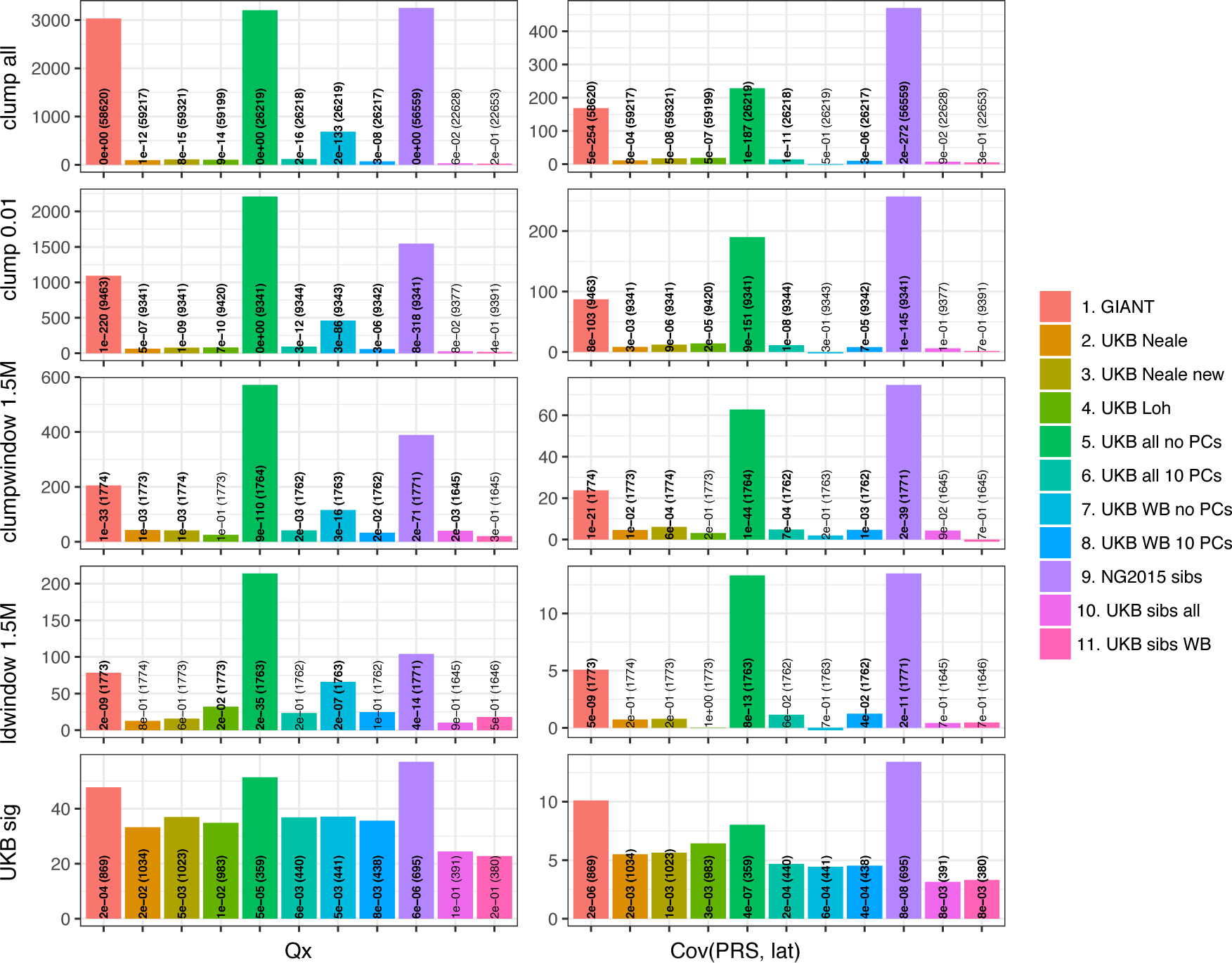
Test statistics for *Q*_*x*_ (left) and latitude correlation ( right) in the POPRES dataset for different summary statistics. The numbers indicate P values and the number of SNPs, and numbers in bold highlight nominal significance (p < 0.05).

**Figure S18.**
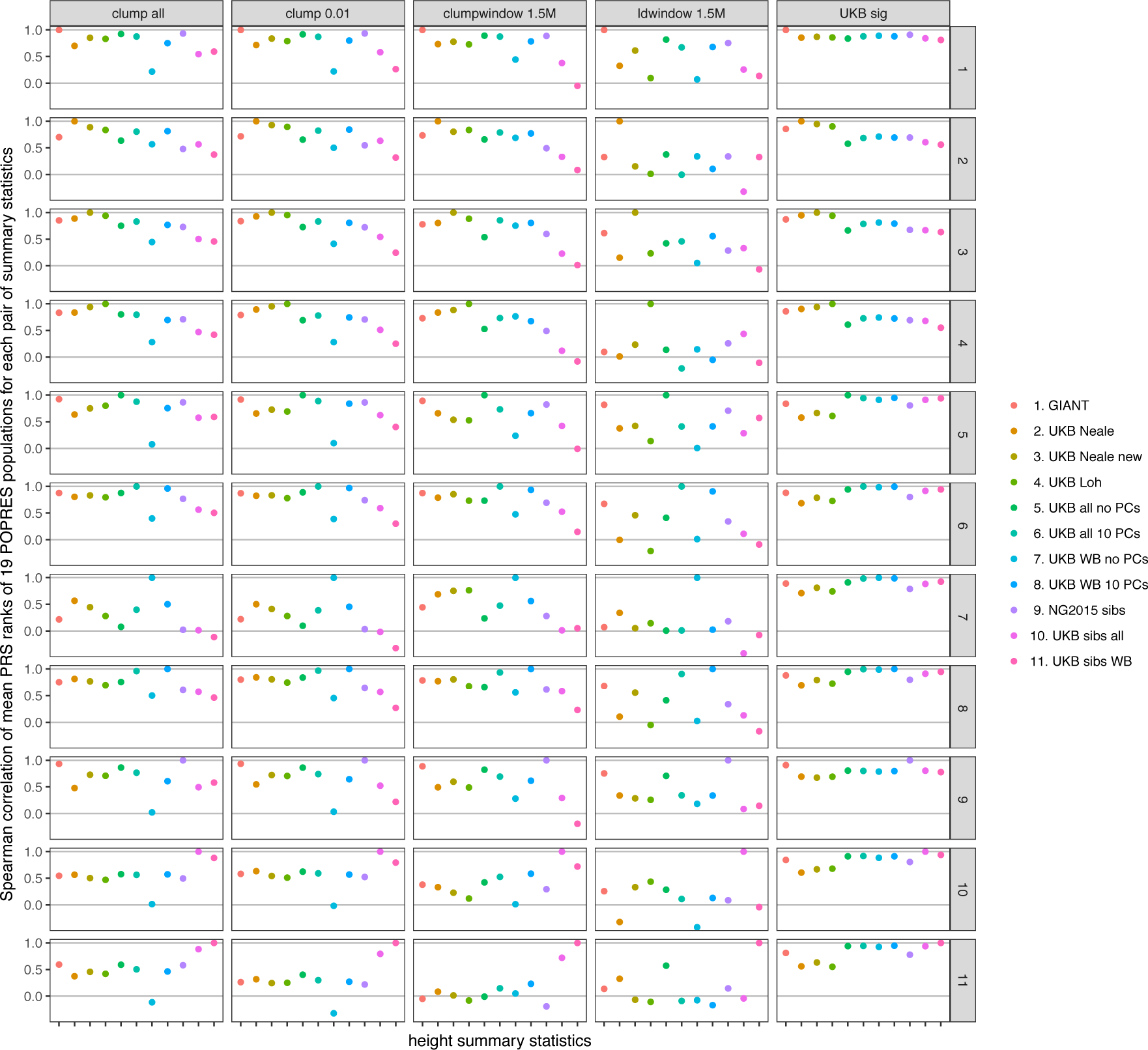
Spearman correlations between polygenic height scores in the POPRES dataset computed from different summary statistics. Spearman correlation coefficients of mean population polygenic score ranking for all pairs of summary statistics at different SNP selections. Polygenic scores from independent SNPs which are genome-wide significant in UKB lead to more consistent rankings than PRS from other sets of SNPs, despite having lower prediction power. Each column shows a different selection of SNPs. clump all: clumped SNPs with no P value threshold; clump 0.01: clumped SNPs with P < 0.01 in UKB and the same number of SNPs in other summary statistics (same as **Figure 4**); clumpwindow 1.5M: genome was split into blocks of 1.5 Mb, lowest P-value SNP was picked in each bin, similar to the 1700 blocks; ldwindow 1.5Mb: genome was split into blocks of 1.5 Mb, random SNP was picked in each bin; UKB sig: LD-pruned SNPs with P < 5×10^−8^ in UKB.

**Figure S19.**
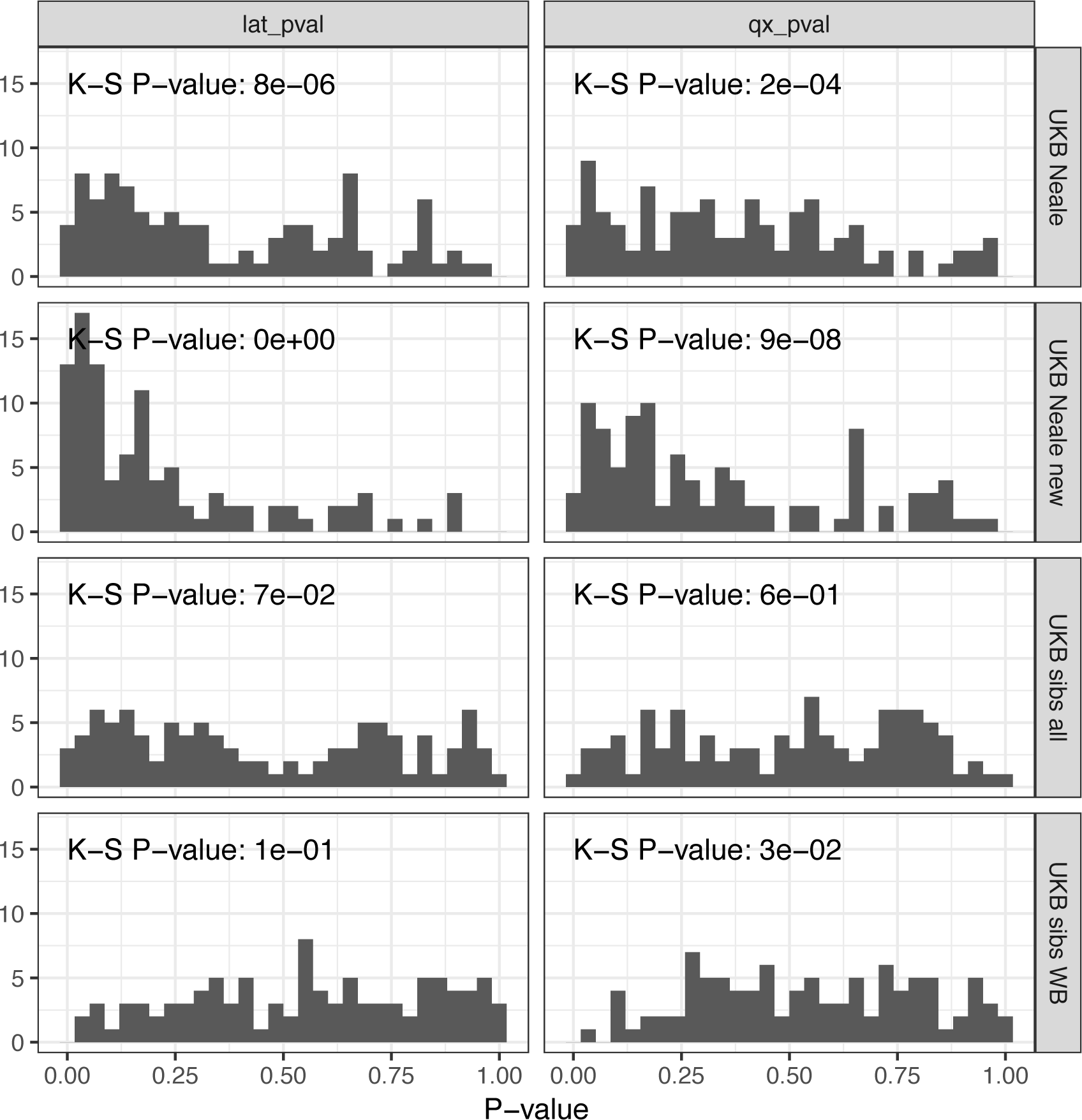
P value calibration in the POPRES dataset for Q_x_ and latitude covariance tests. Random sets of around 1700 independent markers were drawn in 100 repetitions for four summary statistics and Q_x_ and latitude P values were computed. In UK Biobank sibling estimates this resulted in a uniform P value distribution (non-significant Kolmogorov-Smirnov test), while an inflation was observed for UK Biobank GWAS summary statistics.

**Table S1.** Description of 11 GWAS summary statistics.

| name | sample size | number of SNPs (before intersecting) | type | comments | used in figures | PMID | URL |
| --- | --- | --- | --- | --- | --- | --- | --- |
| GIANT | 253288 | 2550858 | GWAS (OLS, meta-analysis) |  | 1, 2, 3, 4 | 25282103 | https://portals.broadinstitute.org/collaboration/giant/images/0/01/GIANT_HEIGHT_Wood_et_al_2014_publicrelease_HapMapCeuFreq.txt.gz |
| UKB Neale (UKB) | 336474 | 10894596 | GWAS (OLS) |  | 1, 2, 3, 4 |  | https://www.dropbox.com/s/sbfgb6qd5i4cxku/50.assoc.tsv.gz |
| UKB Neale new | 360388 | 13791467 | GWAS (OLS) | soon be available on http://www.nealelab.is/ | supp only |  | https://storage.googleapis.com/ukbb-robert/height_ukb_giant/robert1/50.imputed_v3.results.both_sexes.tsv.gz |
| UKB Loh | 458303 | 12007535 | BOLT-LMM |  | supp only | 25642633 | https://data.broadinstitute.org/alkesgroup/UKBB/body_HEIGHTz.sumstats.gz |
| UKB all no PCs | 406825 | 729339 | GWAS (OLS) | genotyped SNPs | 3, 4 |  |  |
| UKB all 10 PCs | 406825 | 729339 | GWAS (OLS) | genotyped SNPs | supp only |  |  |
| UKB WB no PCs | 337208 | 729339 | GWAS (OLS) | genotyped SNPs | 3 |  |  |
| UKB WB 10 PCs | 337208 | 729339 | GWAS (OLS) | genotyped SNPs | supp only |  |  |
| NG2015 sibs | 17500 | 1006257 | within-family (QFAM) |  | 3 | 26366552 | http://cnsgenomics.com/data/robinson_et_al_2015_ng/withinfam_summary_ht_bmi_release_March2016.tar.gz |
| UKB sibs all | 20166 | 729339 | within-family (QFAM) | genotyped SNPs | supp only |  |  |
| UKB sibs WB | 17358 | 729339 | within-family (QFAM) | genotyped SNPs | 3, 4 |  |  |

**Table S2.** Table of ancient and 1000 genomes modern populations used with sample sizes.

| **Population label** | **Population description** | **Meta-population** | **N** |
| --- | --- | --- | --- |
| GBR | British in England & Scotland | Northern Europe | 92 |
| CEU | Utah residents with Northern & Western European Ancestry | Northern Europe | 99 |
| TSI | Toscani in Italia | Southern Europe | 108 |
| IBS | Iberian population in Spain | Southern Europe | 107 |
| PJL | Punjabi from Lahore, Pakistan | South Asia | 96 |
| BEB | Bengali from Bangladesh | South Asia | 86 |
| YRI | Yoruba in Ibadan, Nigeria | Africa | 109 |
| LWK | Luhya in Webuye, Kenya | Africa | 101 |
| JPT | Japanese in Tokyo, Japan | East Asia | 104 |
| CHB | Han Chinese in Beijing, China | East Asia | 103 |
| EF | Ancient | Early farmer | 485 |
| HG | Ancient | Hunter-gatherer | 162 |
| STP | Ancient | Steppe ancestry | 465 |

**Table S3.** Table of POPRES populations used with sample sizes and latitude.

| **Population** | **N** | **Latitude** | **Comments** |
| --- | --- | --- | --- |
| Austria | 14 | 47.5 |  |
| Belgium | 43 | 50.5 |  |
| Czech Republic | 11 | 49.8 |  |
| France | 92 | 46.2 |  |
| Germany | 75 | 51.1 |  |
| Hungary | 19 | 47.1 |  |
| Ireland | 62 | 53.1 |  |
| Italy | 225 | 41.8 |  |
| Netherlands | 17 | 52.1 |  |
| Poland | 22 | 51.9 |  |
| Portugal | 135 | 39.3 |  |
| Romania | 14 | 45.9 |  |
| Spain | 137 | 40.4 |  |
| Sweden | 11 | 60.1 |  |
| Swiss-French | 760 | 46.5 | Lausanne |
| Swiss-German | 84 | 47.3 | Zurich |
| Switzerland | 168 | 46.8 |  |
| United Kingdom | 390 | 55.3 |  |
| Yugoslavia | 44 | 43.9 |  |

**Table S4.** LD Score regression estimates for 11 different summary statistics. LD score regression can be used to detect residual stratification effects in summary statistics, and so we tested whether LDSC confirms our hypothesis of residual stratification. We detect a greatly inflated intercept estimate of 9.42 in UKB all no PCs, but only a moderately increased intercept value in GIANT and an intercept less than one in NG2015 sibs. The relatively small GIANT intercept can be explained by cohort-wise lambda-GC correction, while the low intercept in NG2015 sibs is possibly caused by the adaptive permutation procedure which does not compute precise p-values for non-significant associations. In both cases LDSC cannot be expected to pick up stratification effects, since the generation of summary statistics is not in line with the LDSC model.

| Name | h2 | h2 SE | lambda GC | mean chisq | intercept | intercept SE | ratio | n |
| --- | --- | --- | --- | --- | --- | --- | --- | --- |
| GIANT | 0.31 | 0.01 | 2.00 | 2.92 | 1.28 | 0.02 | 0.15 | 253288 |
| UKB Neale (UKB) | 0.40 | 0.02 | 2.39 | 4.32 | 1.40 | 0.03 | 0.12 | 336474 |
| UKB Neale new | 0.42 | 0.02 | 2.48 | 4.69 | 1.43 | 0.03 | 0.12 | 360388 |
| UKB Loh | 0.60 | 0.03 | 3.31 | 7.66 | 1.75 | 0.04 | 0.11 | 458303 |
| UKB all no PCs | 13.76 | 0.22 | 55.97 | 65.24 | 9.42 | 0.08 | 0.13 | 406825 |
| UKB all 10 PCs | 0.48 | 0.02 | 2.37 | 4.69 | 1.39 | 0.04 | 0.11 | 406825 |
| UKB WB no PCs | 0.54 | 0.03 | 2.67 | 4.66 | 1.66 | 0.04 | 0.18 | 337208 |
| UKB WB 10 PCs | 0.51 | 0.03 | 2.17 | 4.19 | 1.30 | 0.04 | 0.09 | 337208 |
| NG2015 sibs | 0.49 | 0.03 | 1.07 | 1.08 | 0.90 | 0.01 | -1.21 | 17500 |
| UKB sibs all | 0.45 | 0.05 | 0.93 | 1.08 | 0.93 | 0.01 | -0.85 | 20166 |
| UKB sibs WB | 0.47 | 0.06 | 0.93 | 1.06 | 0.93 | 0.01 | -1.18 | 17358 |

**Table S5.** Correlation of beta estimates at all 86,153 shared SNPs.

|  | GIANT | UKB<br>Neale<br>(UKB) | UKB<br>Neale<br>new | UKB<br>Loh | UKB<br>all no<br>PCs | UKB<br>all 10<br>PCs | UKB<br>WB<br>no<br>PCs | UKB<br>WB<br>10<br>PCs | NG2015<br>sibs | UKB<br>sibs<br>all | UKB<br>sibs WB |
| --- | --- | --- | --- | --- | --- | --- | --- | --- | --- | --- | --- |
| GIANT | 1.000 |  |  |  |  |  |  |  |  |  |  |
| UKB<br>Neale<br>(UKB) | 0.660 | 1.000 |  |  |  |  |  |  |  |  |  |
| UKB<br>Neale<br>new | 0.670 | 0.982 | 1.000 |  |  |  |  |  |  |  |  |
| UKB<br>Loh | 0.691 | 0.912 | 0.924 | 1.000 |  |  |  |  |  |  |  |
| UKB all<br>no PCs | 0.332 | 0.276 | 0.274 | 0.251 | 1.000 |  |  |  |  |  |  |
| UKB all<br>10 PCs | 0.673 | 0.973 | 0.971 | 0.916 | 0.290 | 1.000 |  |  |  |  |  |
| UKB<br>WB no<br>PCs | 0.635 | 0.952 | 0.933 | 0.865 | 0.270 | 0.929 | 1.000 |  |  |  |  |
| UKB<br>WB 10<br>PCs | 0.660 | 0.998 | 0.981 | 0.911 | 0.276 | 0.975 | 0.953 | 1.000 |  |  |  |
| NG2015<br>sibs | 0.368 | 0.261 | 0.265 | 0.276 | 0.151 | 0.271 | 0.238 | 0.261 | 1.000 |  |  |
| UKB<br>sibs all | 0.264 | 0.325 | 0.326 | 0.351 | 0.095 | 0.328 | 0.307 | 0.324 | 0.114 | 1.000 |  |
| UKB<br>sibs WB | 0.250 | 0.312 | 0.312 | 0.336 | 0.088 | 0.311 | 0.296 | 0.311 | 0.108 | 0.928 | 1.000 |

**Table S6.**
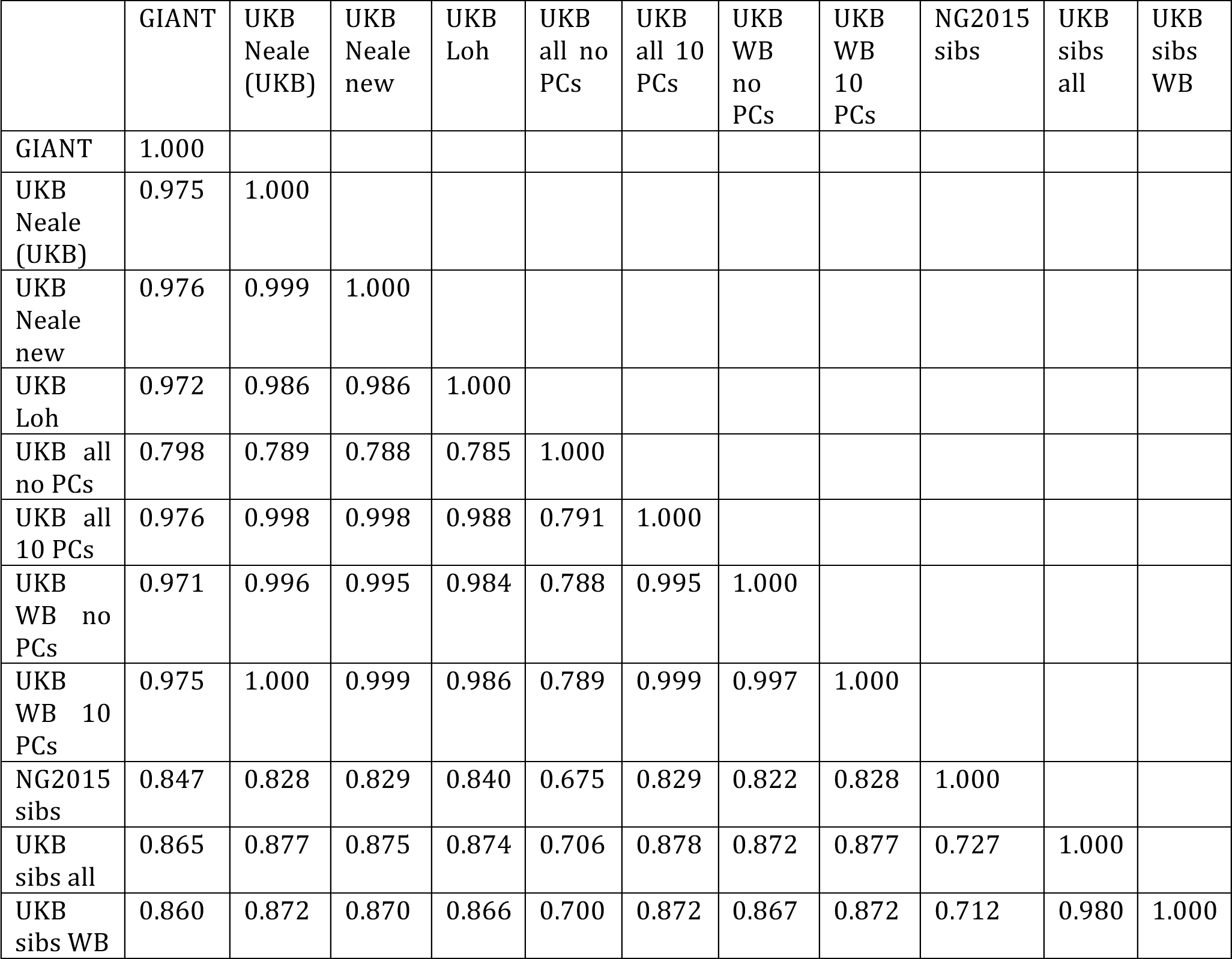
Correlation of beta estimates at 2,251 shared SNPs which are significant in the UK Biobank.

